# Temporal-spatially metabolic reprogramming rewires the H3K27ac landscape to enable the initiation of liver regeneration

**DOI:** 10.1101/2024.07.07.602368

**Authors:** Qi Zheng, Xiaojiaoyang Li, Zhenyu Xu, Yajie Cai, Fanghong Li, Xiaoyong Xue, Shuo Li, Rong Sun, Guifang Fan, Jianzhi Wu, Jiaorong Qu, Runping Liu

## Abstract

The liver possesses extensive regenerative capacity. Nevertheless, the most proximal events driving the transition from quiescent to proliferative hepatocytes remain largely elusive. Using the combination of spatiotemporal metabolomics and transcriptomics, our study mapped out the temporal-spatial landscape of metabolic reprogramming, epigenetic remodeling, and transcriptomic rewiring from 3 to 12 hours post-partial hepatectomy. Specifically, we identified a profound metabolic shift towards hyperactive fatty acid oxidation (FAO) and suppressed phospholipid biosynthesis during the preparation phase of liver regeneration, which were surprisingly reversed afterwards. FAO-dependent accumulation of Acetyl-CoA particularly remodeled H3K27ac landscape. These metabolic reprograming and epigenetic regulation were spatially specific, aligning with the zonation of hepatocyte proliferation. Blocking FAO in etomoxir-treated or hepatocyte-specific *Cpt1a* knockout mice, suppressing Acetyl-CoA biosynthesis, and inhibiting histone acetyltransferase all resulted in lethal liver regeneration deficiency. CUT&Tag analysis further revealed that the reshaping of H3K27ac profiles favored the transcription of genes associated with cell cycle transition and mitosis, and rewired the metabolic gene network. Collectively, we highlight a previously underappreciated role of FAO in epigenetic remodeling that is essential for the initiation of liver regeneration, offering exciting opportunity for the rescue of regeneration-deficient livers.

**Graphical Abstract:** 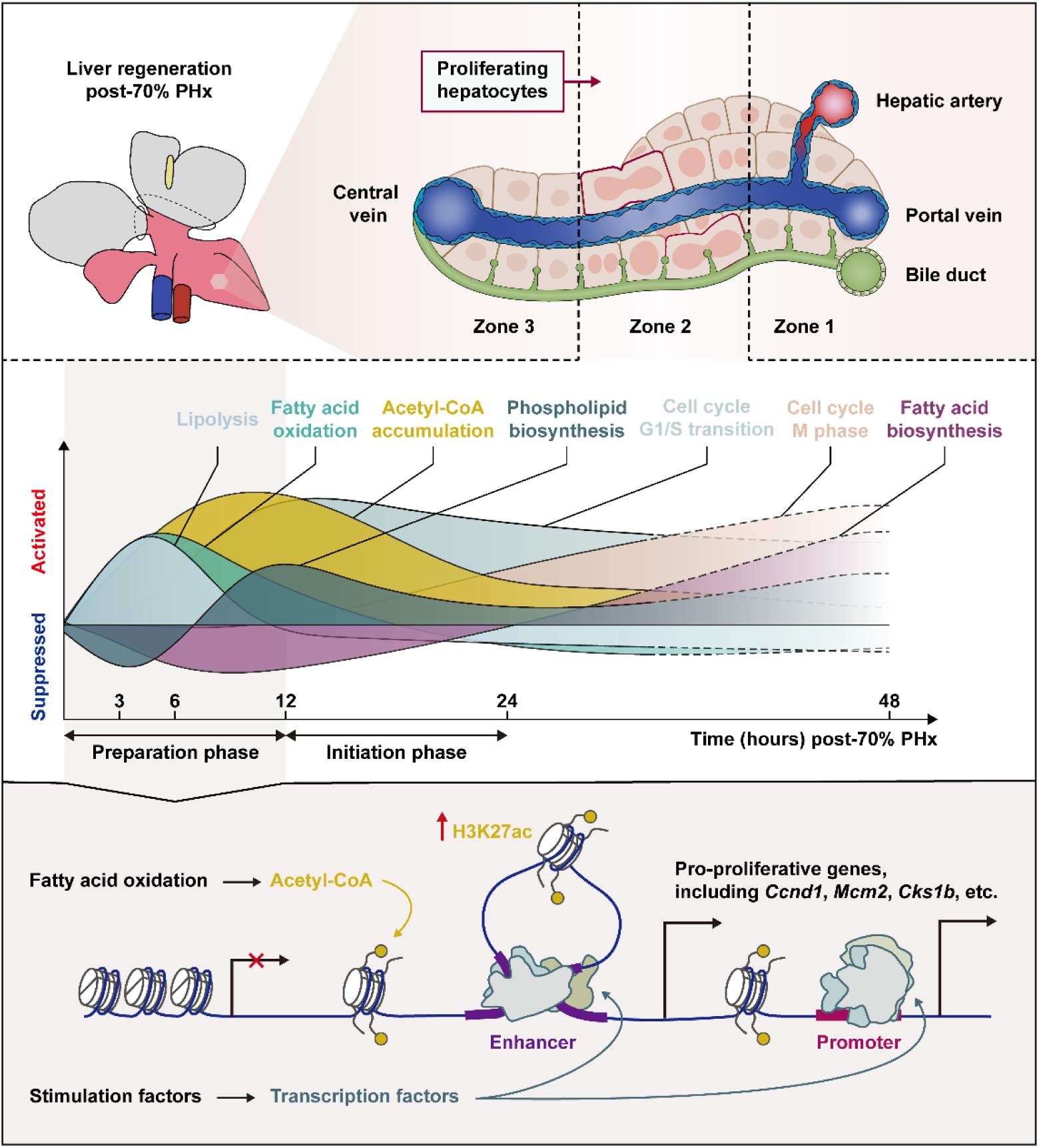

## Introduction

The liver is one of the few organs possessing extensive regenerative capacity, allowing quiescent hepatocytes to rapidly re-enter the cell cycle *in situ* following injuries (*1*). As the incidence of end-stage liver disease continues to rise, the escalating contradiction between limited liver sources and the growing need for liver transplants becomes increasingly pronounced (*2*). Additionally, many patients with insufficient liver functional reserve cannot undergo curative surgeries such as liver resection due to their lack of liver regenerative capacity, or putting them at risk of post-surgery mortality. Deciphering the critical biological events driving the initiation of liver regeneration could lead to advancements in *ex vivo* expansion of hepatocytes as an alternative source for liver transplantation, or improve outcomes for patients undergoing liver resection, opening a new chapter in regenerative medicine for treating liver diseases.

Currently, existing evidence generally supports two major theories regarding the initiation of liver regeneration. The inflammatory response, especially injury-invoked IL-6 and TNF-α signaling has been identified as an important cue to enhance liver regeneration (*3, 4*). The importance of growth factors (e.g., HGF, EGFR, FGF, etc.) and downstream signaling like Hippo and Wnt pathways have also been fully dissected using molecular and genetic approaches (*5, 6*). However, previous studies have primarily focused on the 24 hours to 7 days timeframe following liver injury (*7, 8*), potentially overlooking the poorly-identified preparation and initiation phase of liver regeneration. This might explain why current *ex vivo* simulations based on these theories are ineffective in promoting the re-proliferation of quiescent liver cells. Hence, identifying essential preparatory steps for overriding the blockage between the quiescent state and proliferating phase, particularly within the first 12 hours post-injury, is crucial.

It has been well-established that remnant liver tissue undergoes dynamic metabolic changes in response to partial hepatectomy (PHx) even before the occurrence of increased mitotic activity (*9*). For instance, in rodents, PHx triggered a rapid fall in blood glucose levels at 3 hours post-PHx, followed by a dramatic depletion of glycogen stores during the first 8 hours (*10, 11*). Another well-known characteristic feature of regenerating livers is transient lipid accumulation within hepatocytes preceding the peak proliferative phase, which typically occurred at around 48 hours post-PHx (*12*). Furthermore, due to the exponential cell turnover and the extensive need for cellular membrane biosynthesis, changes in phospholipid homeostasis play a crucial role in the process of liver regeneration and may be associated with the zonation of hepatocyte repopulation (*13, 14*). However, the landscape of spatiotemporally resolved metabolic rewiring in the preparation and initiation phase of liver regeneration remains unclear, presenting both opportunities and challenges.

Metabolic reprogramming has emerged as a major biological node governing the epigenetic landscape by providing substrates and co-factors (*15*). Acetyl-CoA, produced primarily from glycolysis and fatty acid oxidation (FAO), could directly manipulate global histone modification and gene expression. A recent study revealed that FAO abrogation in cardiomyocytes activated the α-ketoglutarate-dependent lysine demethylase, resulting in lowered H3K4me3 peaks in genes necessary for cardiac maturation and promoting cardiomyocyte proliferation (*16*). However, several other lines of evidence suggest that the overexpression of ACLY and ACSS2, by enhancing Acetyl-CoA flux, are pertinent to support the regeneration of various tissues, including skeletal muscle, bone marrow, and hippocampal neuron (*17–20*). Moreover, the epigenetic remodeling has been found to license adult liver cells for hepatic organoids formation and liver regeneration (*21, 22*). One of the earliest studies also found a significant temporary increase in histone acetyltransferase (HAT) activity before the onset of DNA replication in regenerating liver (*23*). These findings collectively suggest a possibility that links metabolic reprogramming with epigenetic modification-driven liver regeneration. However, whether this connection is a prerequisite for the exit of hepatocytes from quiescence and the spatial-temporal characteristics of this connection warrant further investigation.

In this study, we utilized integrated transcriptomics, metabolomics, and spatial metabolomics to delineate the spatiotemporal landscape of metabolic reprogramming within the first 12 hours following liver resection. Additionally, leveraging CUT&Tag analysis, we elucidated the changes in the histone acetylation landscape driven by FAO through increased Acetyl-CoA supply, thereby establishing a network that interconnects metabolic reprogramming, epigenetic regulation, and transcriptomic reshaping during the preparation and initiation of liver regeneration.

## Materials and methods

### Animal studies

All animals received human care, mouse experiments were performed in strict accordance with guidelines of Chinese Association for Laboratory Animal Science and all other relevant national and institutional guidelines, and approved by the Institutional Animal Care and Use Committee of Beijing University of Chinese Medicine (BUCM-2023082201-3058).

Male C57BL/6J mice (6-8 weeks old) were purchased from SPF Biotechnology Co., Ltd. (Beijing, China). *Cpt1a*^fl/fl^ mice (*Cpt1a*^fl/fl^ allele) were generated by inserting loxP sites within intron 1 and intron 2, flanking exon 2 of *Cpt1a*, with assistance from Gempharmatech CO., Ltd. in Nanjing, China. Inducible hepatocyte-specific CPT1A knockout mice (LKO) were established *via* the tail vein injection of HBAAV2/9-TBG-cre-ZsGreen viruses (500 μL, 1.8*10^12^ viral genomes/mL) with HBAAV2/9-TBG-ZsGreen viruses (500 μL, 1.9*10^12^ viral genomes/mL) as controls (obtained from Hanbio Biotechnology Co., Ltd., Shanghai, China). After 3 weeks of infection, the GFP expression in mice liver was observed using fluorescence microscopy. All mice were housed in a barrier facility under specific pathogen-free conditions, at 22 ± 1 °C and a relative humidity of 50 ± 5% with controlled illumination (12 h dark/light cycle). Food and tap water were available *ad libitum* during the whole experiment.

### Partial hepatectomy and sample preparation

Mice were anaesthetized using 3-4% isoflurane and maintained on 2.5% isoflurane throughout the surgery, meanwhile the core body temperature was maintained at 37.5 °C using a heated platform. To build 70% PHx-induced liver regeneration models, the left lobe and the median lobe were ligated separately and resected, as described (*24*). As control, sham operations were performed but livers were not removed. For time-course analysis of early liver regeneration, mice were randomized into different groups and sacrificed at 3 hours, 6 hours, 12 hours, 24 hours, and 48 hours post-PHx, respectively. Liver samples were harvested and immediately snap-frozen in liquid nitrogen for the following experiments. For immunohistochemistry analysis, a portion of the caudate lobe was excised and fixed in 4% paraformaldehyde solution for further embedding in paraffin. For spatiotemporal metabolome analysis, a portion of the right lateral lobe was embedded in Cryo-Gel (Leica, Germany) and were stored at −80 °C until cryo-sectioning.

### Inhibitors treatment

In the subsequent *in vivo* experiments, etomoxir (CPT1A inhibitor, 12.5 mg/kg), 2-deoxy-D-glucose (HK2 inhibitor, 500 mg/kg), sodium oxamate (LDHA inhibitor, 500 mg/kg), ACSS2i (ACSS2 inhibitor, 25 mg/kg), or C646 (P300 inhibitor, 15 mg/kg) were administered by intraperitoneal injection, while 10,12-Tricosadiynoic acid (ACOX1 inhibitor, 100 mg/kg) or ETC-1002 (ACLY inhibitor, 30 mg/kg) was given by gavage. Sham control and solvent control were included in all assays.

### Quantitative reverse transcription PCR

Total RNA was extracted from snap frozen liver tissues using TRIzol reagent. 1 μg of total RNA was reverse transcribed to cDNA by HiScript III RT SuperMix for qPCR kit. qRT-PCR was performed on Bio-Rad CFX-96 Touch system (CA, USA) using Taq Pro Universal SYBR qPCR Master Mix. Primer sequences are shown in detail in the **Supplementary information**. The expression of genes was normalized to *Hprt1* levels and were shown in the figures. All reagents were purchased from Vazyme Biotech (Nanjing, China).

### Histology and immunohistochemistry

Histology and immunohistochemistry were performed using standard protocols. For descriptions of protocols and antibodies used, see the **Supplementary information**.

### Western blotting analysis

Protein sample preparation and Western blotting were performed using standard techniques. For descriptions of protocols and antibodies used, see the **Supplementary information**.

### RNA sequencing

Approximately 1 μg of total RNA extracted from mouse livers (n=3 or 4 for each group) was used for RNA-seq library preparation following the Illumina TruSeq stranded mRNA sample preparation guide (Illumina, San Diego, CA). The methods for RNA sequencing and analysis were described in **Supplementary information**.

### Untargeted Metabolomics

LC-MS/MS analysis-based untargeted metabolomics was carried out on the mice liver samples (n=4 to 6 for each group). LC-MS/MS analysis was performed using a Vanquish UHPLC system (Thermo Fisher, Germany) coupled with an Orbitrap Q Exactive^TM^ HF-X mass spectrometer (Thermo Fisher, Germany). Details about untargeted metabolomics detection and data analysis were shown in **Supplementary information**.

### Airflow-assisted desorption electrospray ionization mass spectrometry imaging analysis

Frozen mouse liver tissue samples (n=3 for each group) from the right lateral liver lobe were fixed with Cryo-Gel medium. 3 adjacent slices were prepared, one for H&E staining and the other for AFADESI-MSI image. The AFADESI-MSI images were superimposed with the H&E-stained images. Details of AFADESI-MSI analysis were given in the **Supplementary information**.

### CUT&Tag analysis

For the CUT&Tag assay, we followed the CUT&Tag-direct protocol described by Henikoff *et al*. (*25*). Bulk H3K27ac CUT&Tag libraries generated from liver samples were used as a mark for transcriptional activation. Details of CUT&Tag analysis were given in the **Supplementary information**.

### Binding and Expression Target Analysis for integrating CUT&Tag and RNA-seq results

BETA was performed to predict whether H3K27ac has activating or repressive function by combining CUT&Tag and RNA-seq results. The analysis pipeline was performed as previously described (*26*). Briefly, BETA estimates H3K27ac’s regulatory potential score for each gene based on the distance between H3K27ac binding sites and TSSs of each gene, and also based on the number of H3K27ac binding sites ± 50 kb centered at the TSS of each gene. BETA then uses a nonparametric statistical test (Kolmogorov-Smirnov test) to compare regulatory potential scores for genes that are up-target, down-target or not regulated based on RNA-seq results with and without H3K27ac overexpression.

### Statistical analysis

For all quantitative analyses, results are representative of at least three independent experiments or at least eight mice for each group. Data were expressed as mean ± SEM and statistical analysis was performed using Prism 8.0.2 (GraphPad, La Jolla, CA, USA). Statistical tests were used based on the assumption that sample data are derived from a population following a probability distribution based on a fixed set of parameters. Two-tailed Student’s T-test was performed to determine the statistical significance of differences between two groups. One-way ANOVA was used for multiple comparison tests. Survival curve analysis was performed using Kaplan-Meier survival analysis. Log-rank (Mantel-Cox) test was used for the survival curve comparison, and 95% CI from the Mantel-Haenszel test were used for the hazard ratios. *P*<0.05 were considered statistically significant (\**P*<0.05; \*\**P*<0.01; \*\*\**P*<0.001; \*\*\*\**P*<0.0001; *P*>0.05, n.s. = not significant).

## Results

### The first 12 hours post 70% partial hepatectomy is the preparatory phase of liver regeneration

To investigate the dynamics of hepatocyte proliferation during liver regeneration, we obtained liver tissues from mice at various time intervals, especially within the first 12 hours, following 70% PHx (**Fig. 1A**). Analysis of regenerating liver tissue through hematoxylin and eosin (H&E) staining did not reveal any abnormal hepatic lesions such as inflammation, fibrosis, or necrosis (**Fig. S1A**). Regenerated hepatocytes were observed in the livers at 24 and 48 hours post-PHx, as revealed by KI67 and PCNA immunohistochemical staining (**Fig. 1B-C**), as well as qPCR results (**Fig. 1D**). Remarkably, we observed the re-entry of the cell cycle in hepatocytes at earlier time points, as demonstrated by the persistent upregulation of cell cycle progression marker *Ccnd1* from as early as 6 hours post-PHx.

**Fig. 1.**
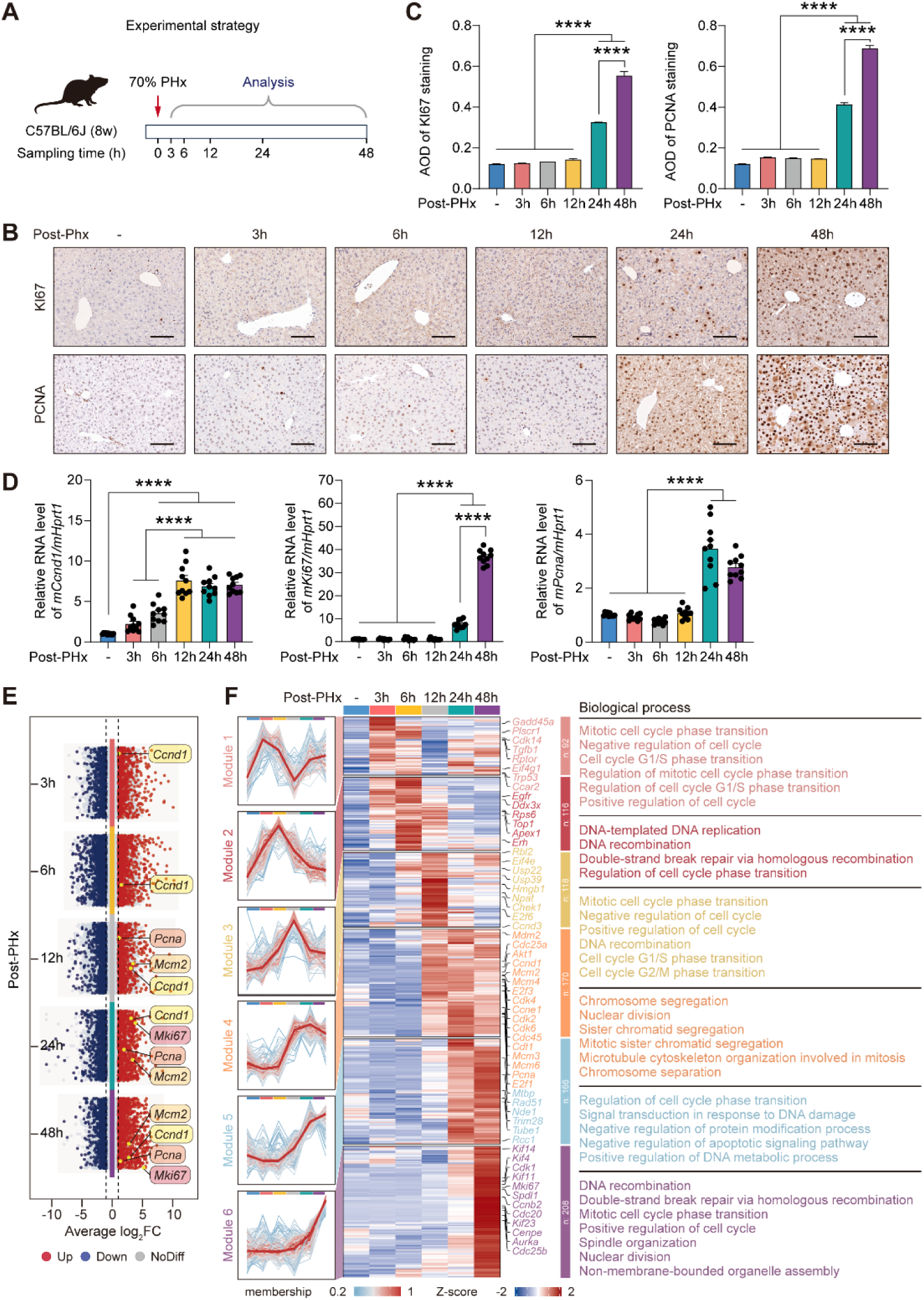
Time-course analysis of proliferative phenotype during the early phase of liver regeneration following 70% PHx. (A) Experimental design. (B) Immunohistochemical results for KI67 and PCNA in regenerating livers at different times following the surgery. Scale bars, 100 μm. AOD, average optical density. Quantifications were performed on at least 5 different areas of the sections in a random way. (C) The average optical density of KI67 and PCNA staining. (D) qRT-PCR analysis of *Ccnd1*, *Ki67* and *Pcna* expressions in mouse livers at different regenerating stages. Data are represented as mean ± SEM of 10 replicates. (E) Multiple volcano plot of differential gene expression analysis of regenerating livers at different times compared to control. Red/blue symbols classify the upregulated/downregulated genes according to the criteria: |log_2_FC| > 1 and adjusted *P*-value < 0.05. Dashed lines represent the average log_2_FC value +1 and −1, respectively. (F) Six gene modules were identified to reveal time-dependent proliferation-related transcriptome specificity using HCA. Middle panel: The heatmap represents the expression of proliferating genes involved in different phase of early liver regeneration within 48 hours post-PHx. Left panel: The coexpression patterns of the hierarchical clustered genes in the six modules. Right panel: GOBP enrichment for each gene module. \**P* < 0.05, \*\**P* < 0.01, \*\*\**P* < 0.001, \*\*\*\**P* < 0.0001, One-way ANOVA. Error bars represent mean ± SEM.

As shown in **Fig. S1B**, significant transcriptomic reshaping was observed as early as 3 hours post-PHx (**Fig. 1E**). The hierarchical clustering analysis (HCA) and gene ontology biological processes (GOBP) enrichment analysis identified six gene modules characterizing the sequential activation of liver regeneration (**Fig. 1F**). Specifically, module 1 and 2 represent genes highly expressed at 3 and 6 hours after PHx, respectively. These differentially expressed genes (DEGs) were associated with DNA replication and G1/S transition, suggesting the re-entry of quiescent hepatocytes into the cell cycle. DEGs in Module 3, with a sharp upregulation at 12 hours post-PHx and a gradual decrease later on, were involved in G1/S or G2/M cell cycle transition. DEGs in Module 4, with increased abundances persistently from 12 to 48 hours after PHx, were enriched in biological processes such as chromosome segregation, nuclear division, and microtubule cytoskeleton organization, which are all involved in mitosis. As liver regeneration speeds up after 24 hours post-PHx, the upregulated genes related with homologous recombination DNA repair, organization of the microtubules, spindle assembly, and cytokinesis dynamics, were highly expressed, showing a positive correlation with *Ki67* expression. These results depict significant temporal changes in the hepatic transcriptome, emphasizing the importance of focusing on the first 3 to 12 hours following 70% liver resection for understanding key biological processes driving the initiation of liver regeneration.

### Integrative transcriptomics and temporal-spatially metabolomics delineates dynamic metabolic reprograming during liver regeneration

Metabolic reprograming has been considered as a hallmark of liver tissue self-renewal and adaptation (*12*). Accordingly, HCA based on RNA-seq datasets revealed a significant alteration in the transcriptional profiles of genes involved in energy metabolism in the initial 12 hours following PHx (**Fig. 2A**). The expression of DEGs related to pyruvate metabolic process, glycogen catabolic process, polysaccharide metabolic process, and glycolytic process showed the most drastic and continuous decrease throughout the initial 48 hours post-PHx. In contrast, the expression of DEGs in Module 2 and Module 3 were dramatically upregulated as early as 3 to 6 hours after liver resection, long before the entry of hepatocytes into mitosis. These genes were enriched in pathways like lipid catabolic process, fatty acid β-oxidation, acyl-CoA metabolism, carnitine metabolism, tricarboxylic acid cycle, and aerobic respiration, thus generally contributing to the activation of FAO. The transient increase in the expression of genes involved in glycolysis, and the biosynthesis of key metabolites, such as phospholipid, steroid, and glycogen, were mostly reversed after 12 hours post-PHx (Module 4). In addition to these processes, DEGs in Module 5 were further enriched in pathways related to fatty acid and triglyceride biosynthesis, complementing the temporal gene expression patterns involved in lipid catabolism and FAO. These findings represent a significant shift of hepatic metabolism towards lipid degradation during the preparatory stage of liver regeneration.

**Fig. 2.**
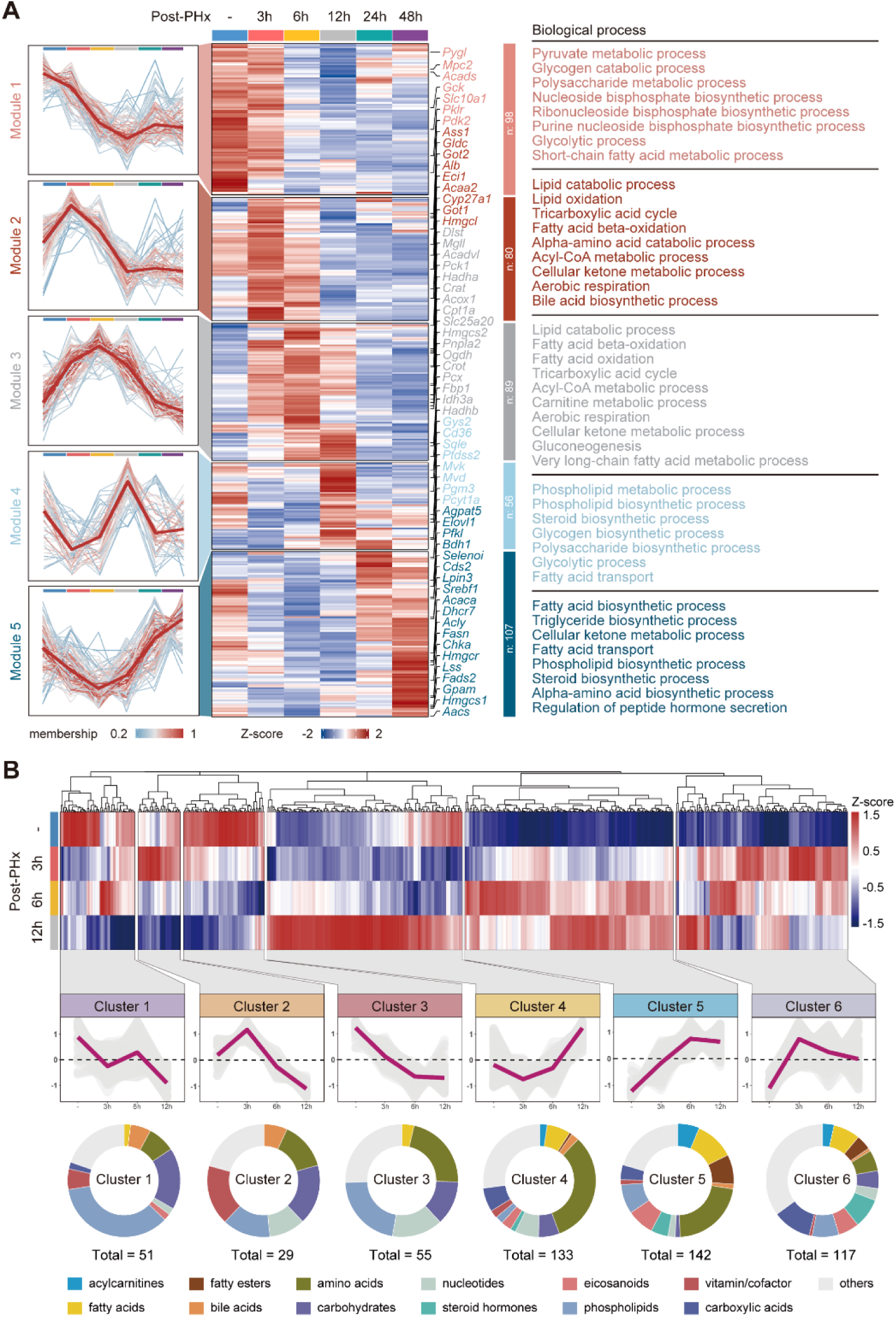
Rewired metabolic phenotype is a hallmark of regenerating liver. (A) Time-dependent metabolism-related transcriptome specificity in five gene modules. Left panel: HCA for different clusters. Middle panel: The heatmap of HCA visualizes the expression of total metabolic genes within 48 hours post-PHx. The clusters of genes associated with similar biological processes are grouped based on the degree of GOBP enrichment, as indicated on the right side. (B) The heatmap represents the untargeted metabolomics landscape across liver samples within the priming phase of regeneration. The pie-chart shows the relative contribution of metabolites features in different clusters.

To further validate the metabolic phenotype conveyed by the transcriptomic changes, untargeted metabolomics was employed to depict a time-resolved metabolic landscape during the early stages of liver regeneration. Principal component analysis and volcano plots indicated significant changes in the intrahepatic metabolic profiles at different time points after PHx (**Fig. S1C and D**). HCA further sub-divided all differential metabolites into 6 clusters based on alteration patterns during the initial 12 hours of liver regeneration (**Fig. 2B**). A significant downregulation over time was observed in cluster 1 to 3, which was characterized by a simultaneous decrease in levels of phospholipids, vitamins/cofactors, and carbohydrates. On the other hand, cluster 4-6 exhibited significant accumulations of other metabolites from 3 to 12 hours post-PHx, such as acylcarnitines (ACars), fatty acids, fatty esters, eicosanoids, and steroid hormones. The levels of various ACars, which are essential intermediate metabolites in FAO, continued to increase until 48 hours after PHx (**Fig. S1E**). Taken together, these results provide additional insights into the metabolic reprogramming post-PHx, primarily involving activated FAO, Acetyl-CoA accumulation, and suppressed biosynthesis of phospholipid, and fatty acid, emphasizing the strong connection between the persistent activation of FAO and the initiation of liver regeneration.

### Spatially specific activation of lipid catabolism aligns with the zonation of liver regeneration

The liver exhibits spatial “zonation”, with distinct metabolic functions and secretion allocated along the lobular axis of blood flow. This hierarchical structure divides the liver lobule into three zones - Zone 1 (portal vein areas), Zone 3 (central vein areas), and Zone 2 (middle parts), each with distinct roles during liver regeneration. To comprehend the spatial patterns of intrahepatic metabolic reprogramming during early liver regeneration, the right lateral liver lobe from mice sacrificed at 0, 12, or 24 hours after PHx were subjected to AFADESI-MSI based spatially resolved metabolomics analysis as described in the **Methods** and **Fig. S2A**. The t-SNE dimension reduction and cluster analysis based on *in situ* metabolic signatures were performed, showing considerable heterogeneity among the characteristic substances, especially the lipids (**Fig. S2B and C**).

A spatial-temporal metabolome-transcriptome association network was then established for triglyceride lipolysis and FAO pathways. As shown in **Fig. 3A-C**, the upregulation of genes involved in the lipolysis of triglycerides (TG), which occurred as early as 3 to 6 hours post-PHx, was consistent with the accumulation of DG, MG, and fatty acids in the liver, 12 hours post-PHx (**Fig. S2D**). Paradoxically, the intrahepatic levels of certain TGs were also increased, indicating net influx of lipids in the regenerating livers, although few lipids transportation-related genes were differentially expressed. From a spatial perspective, fatty acylglycerols and corresponding FAs, such as TG(34:0), DG(34:1), MG(15:0), FA(18:1), and FA(16:0), are primarily found in Zone 2 and Zone 3 hepatocytes during the resting state. Following the initiation of liver regeneration, particularly 12 hours after liver resection, the levels of these substances universally increased throughout the liver, with the most significant increase observed in Zone 1. As liver regeneration progresses into the rapid proliferation phase, the levels of fatty acylglycerols decreased and tended to the spatial distribution of the resting state, while the levels of FAs remain elevated. The expanded intrahepatic FA pool was accompanied with increased mitochondrial and peroxisomal FAO, as characterized by the synchronous upregulation of CPT1A and ACOX1, and the highly expressed acyl-CoA transporters (SLC25A20, CRAT, and CROT) and dehydrogenases (ACADVL, ACADL, ACADM, and ACADS) (**Fig. 3A and B**). The elevation of FAO products like Acar(18:1) and Acar(16:0), especially their specific enrichment in Zone 2 hepatocytes, represented the predominant response of most ACar substances to the initiation of liver regeneration (**Fig. 3D and E**).

**Fig. 3.**
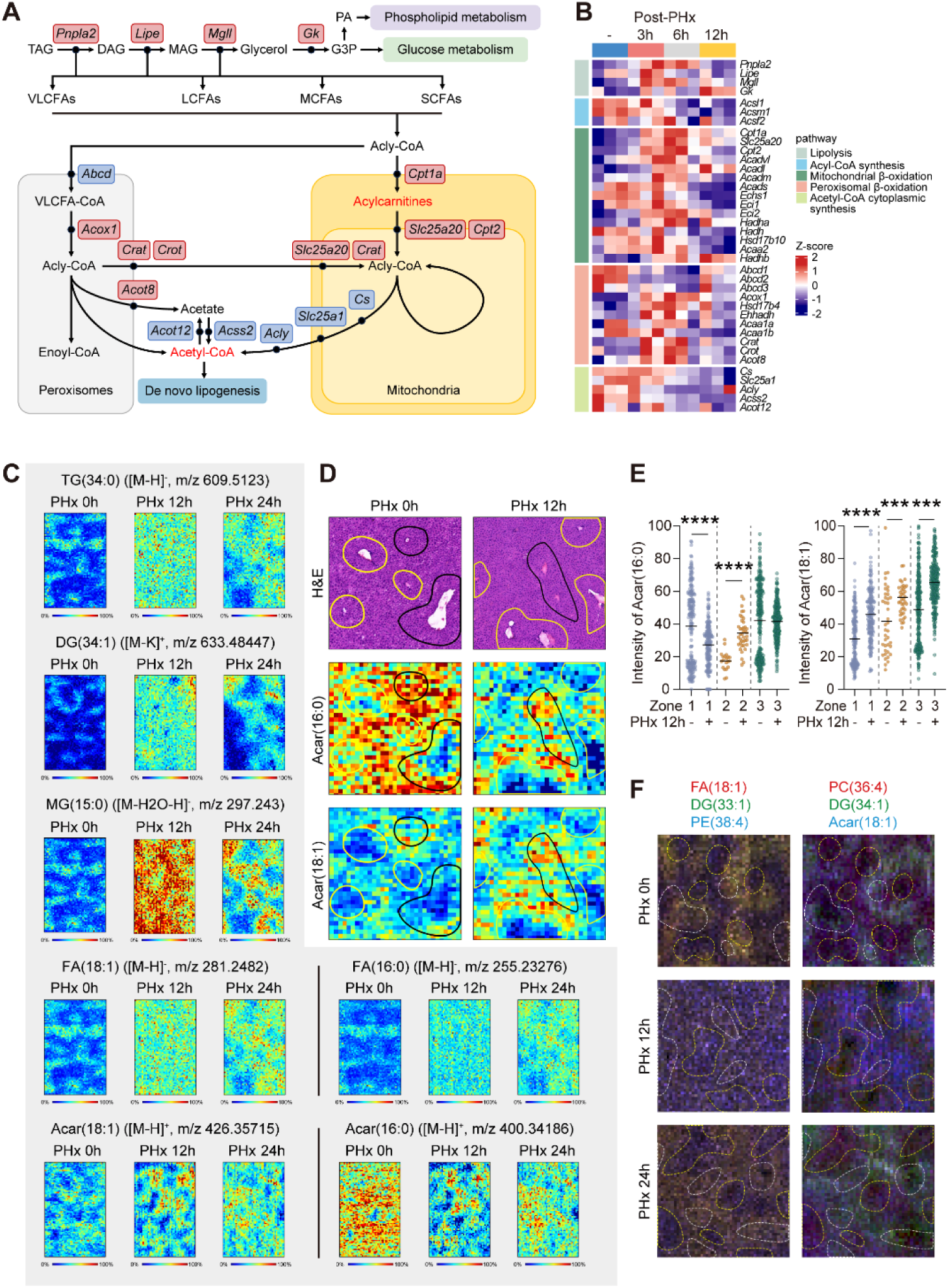
The spatiotemporal evolution of enhanced lipid catabolism in the regenerating liver. (A) Gene expression and metabolic fluxes related to lipolysis and FAO pathways. For triglyceride lipolysis, genes involved in glyceride hydrolysis (*Pnpla2*, *Lipe*, *Mgll*, and *GK*), fatty acid activation (for example, ACS family), carnitine shuttle (*Cpt1a*, *Slc25a20*, and *Cpt2*), mitochondrial oxidation (for example, ACAD family) and peroxisomal oxidation (for example, ACOX family). Acetyl-CoA, generated from FAO, can enter the TCA cycle, producing citrate for *de novo* lipogenesis. Predicted metabolic flux within 12 hours regeneration is indicated by the color of enzyme names (red = upregulated; blue = downregulated). (B) Individual gene expressions after PHx are shown as heatmaps, with legend indicating different metabolic fluxes on the right. (C) MSI images of representative lipids in liver tissues at 12 and 24 hours post-PHx, with color scale intensity representing relative values. (D and E) Enlarged MSI images and levels of Acars in different liver zones. Error bars represent mean ± SEM. \*\*\**P*<0.001; \*\*\*\**P*<0.0001, unpaired Student’s t test. (F) The spatial distribution of FA(18:1), DG(33:1), PE(38:4), PC(36:4), DG(34:1), and Acar(18:1) in control liver is compared to that following PHx.

Robustly inhibited fatty acid biosynthesis (FAB) pathways, including *de novo* synthesis, desaturation, and elongation, further highlighted the reprogramming of hepatic FA metabolism during the initiation of liver regeneration, shifting from balanced synthesis and degradation to predominant catabolism (**Fig. S3A**). Besides, phospholipid metabolism plays a crucial role in lipid homeostasis by serving as a shunt pathway and competing with FA metabolism. Both transcriptomic and metabolomic analysis revealed a perturbated phospholipid metabolism 12 hours post-PHx, leading to an overall reduced phospholipids pool, which were subsequently recovered later on (**Fig. 2B** and **Fig. S3A and B**). Interestingly, the co-localization of representative lipid metabolites further supports the spatial specificity of the metabolic reprogramming towards a bias in FAO (**Fig. 3F**). Within the initial 12 hours of liver regeneration, there was a minimal increase in the overall levels of DG and FA in Zone 2 relative to Zone 1 and Zone 3. Conversely, a noticeable decrease of phospholipids specifically in Zone 2 was observed. These results suggest that in Zone 2, the DG and FA provided by external transport and upregulated lipolysis processes were rapidly depleted by hyperactive FAO, leading to the specific accumulation of ACars in Zone 2. At 24 hours post-PHx, we observed a re-enrichment of DG and FA, an overall increase in phospholipid levels, and continued active FAO in Zone 2. These findings suggest that in response to the metabolic stress following liver resection, or in the preparation for liver regeneration, hepatic lipid metabolism is reprogrammed to favor the dominant FAO pathway and suppress other pathways, spatially overlapping with the zonation of liver regeneration.

### FAO is essential for the initiation of liver regeneration by facilitating H3K27 acetylation

To further explore the biological significance of the spatiotemporal overlap between highly active metabolic reprogramming and hepatocyte repopulation, we performed a correlation analysis between differential metabolite levels and the expression of cell cycle- and proliferation-related genes. In the first 12 hours after liver resection, there was an inverse correlation between the cell cycle progression and metabolites like carbohydrates, PC, and PE. However, the ACar levels were positively related to hepatocyte proliferation (**Fig. 4A**), which was equally significant within 24 to 48 hours post-PHx (**Fig. S4A**). Regression analysis also proved strong positive correlations between ACar levels and the expression of *Ccnd1* and *Ki67* (**Fig. S4B**).

**Fig. 4.**
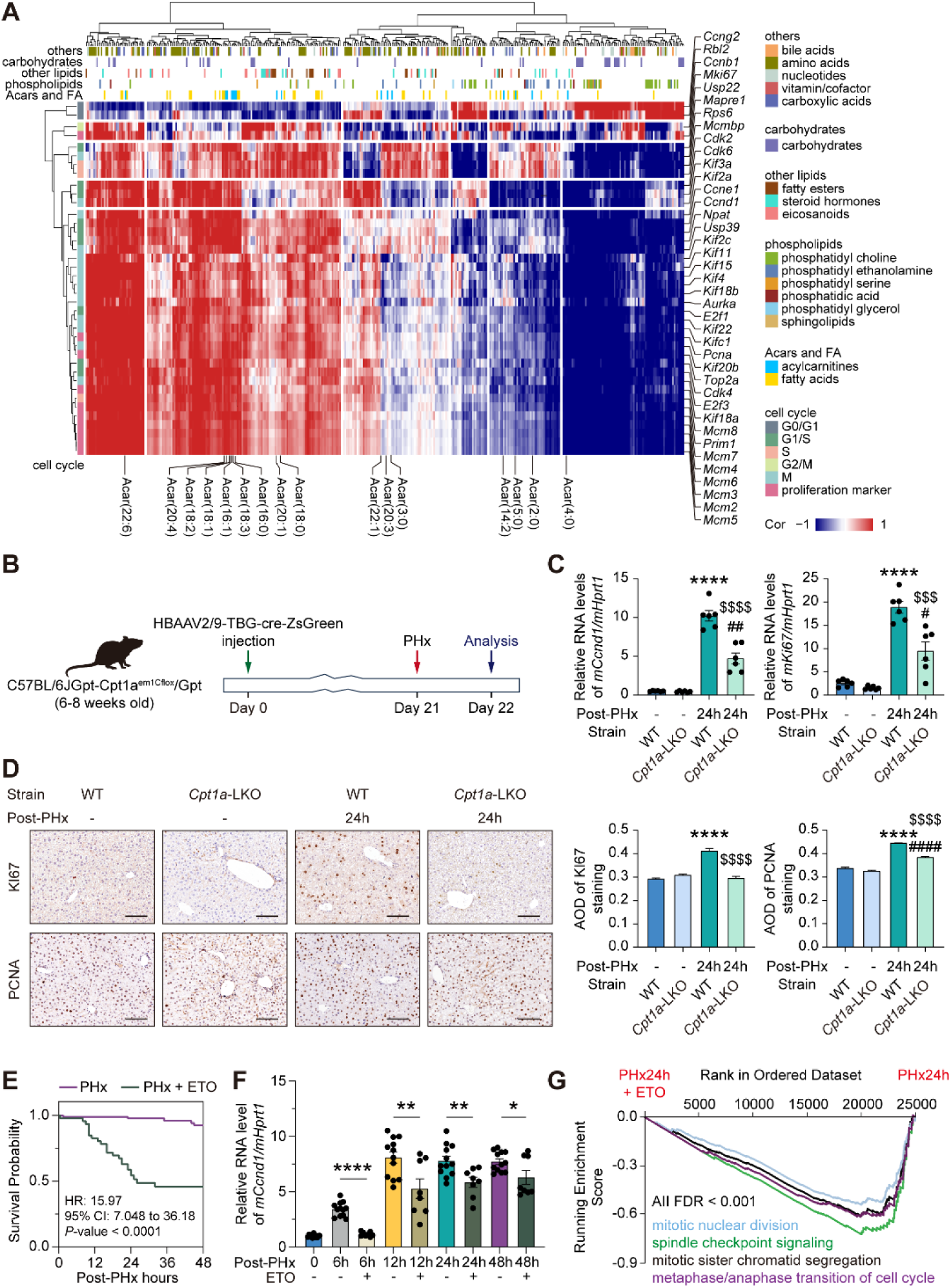
Inhibition of mitochondrial FAO impedes the initiation of liver regeneration. (A) Correlation analysis of the association between PHx-altered proliferating genes and differential metabolites at 12 hours after liver section. The Pearson’s correlation (R) values are visualized in a heatmap. The different types of metabolites analyzed are annotated above the heatmap with details shown in the legend to the right. The color code representing each cell cycle phase is shown in the lower right legend of the heatmap. (B) Schema describing approach to establish hepatocyte specific Cpt1a KO mice. (C) The transcription levels of *Ccnd1* and *Ki67* in LKO mice were determined by qRT-PCR assay. Data are represented as mean ± SEM of 6 replicates. (D) Immunohistochemical staining determines the expression of KI67 and PCNA, with quantitative analysis of AOD value. Quantifications were performed on at least 5 different areas of the sections in a random way. (E) Kaplan-Meier post-PHx survival curve for mice treated with ETO (n=46) or without ETO (n=80). HR, Hazard Ratio. (F) The transcription levels of *Ccnd1* after ETO treatment were determined by qRT-PCR assay. Data are represented as mean ± SEM of 8 or 12 replicates. (G) GSEA enrichment plots showing a significant enrichment (FDR < 0.05) of down-regulated genes (PHx24h+ETO *vs.* PHx24h) in proliferation related pathways. FDR, false discovery rate. \**P*<0.05; \*\**P*<0.01; \*\*\**P*<0.001; \*\*\*\**P*<0.0001, unpaired Student’s t test. Error bars represent mean ± SEM.

Hepatocyte-specific *Cpt1a* knockout (*Cpt1a*-LKO) mice were than established to identify whether mitochondrial FAO is essential for liver regeneration. Briefly, *Cpt1a*^fl/fl^ mice were subjected to AAV-TBG-Cre infection as described in the **Methods** (**Fig. 4B**). The expression of zsGreen in hepatocytes, reduced Cpt1a mRNA levels and significant hepatic steatosis were found in *Cpt1a*-LKO mice (**Fig. S5A-C**). As shown in **Fig. 4D and E**, the expression of CCND1 and KI67 was significantly decreased in these mice, indicating that hepatic deletion of *Cpt1a*impaired PHx-induced liver regeneration. The specific inhibitor of CPT1α, etomoxir (ETO), was further employed to verify the results derived from the transgenic mice (**Fig. S6A**). As depicted in **Fig. S6D**, ETO treatment also resulted in FAO deficiency, leading to lipid accumulation in hepatocytes. This FAO defect significantly increased the mortality of PHx mice post-operation, with the survival rate decreased to 54% within 24 hours (**Fig. 4E**). The higher mortality was attributed to the severe delay in liver regeneration initiation, as indicated by the ineffective induction of *Ccnd1* expression within 6-12 hours post-PHx (**Fig. 4F**). Additionally, FAO deficiency led to a broad suppression of gene expression associated with cell cycle progression and mitotic processes within 24 and 48 hours post-PHx, ultimately reflected in the reduction of KI67- and PCNA-positive cells in the liver (**Fig. 4G** and **Fig. S6B, C, E, and F**). Moreover, it is noteworthy that survivors of ETO treatment were compelled to seek other alternative routes (i.e., aerobic metabolism) to sustain liver reprogramming and survival after PHx (**Fig. S6G**). On the other hand, the inhibition of peroxisomal FAO by TDYA showed minimal effects on the expression of *Ccnd1* (**Fig. S7D and E**). Furthermore, although the rapid consumption of glucose and the reprogramming of glucose metabolism from oxidative phosphorylation to glycolysis were observed (**Fig. S7A-C**), the inhibition of glucose phosphorylation or lactate oxidation in glycolysis by 2-DG and OXA, respectively, had no effect on the initiation of liver regeneration (**Fig. S7D and E**). These results collectively indicate that hyperactive mitochondrial FAO but not other reprogrammed metabolic pathways is essential for the initiation of liver regeneration.

### FAO-derived Acetyl-CoA facilitates H3K27Ac remodeling to enable liver regeneration

The highly active FAO process generates a substantial amount of Acetyl-CoA, which in turn promotes the acetylation of related proteins by providing sufficient substrates. Although the mRNA levels of ACLY and ACSS2 are decreased during liver regeneration, implying deficient intracellular transferring of Acetyl-CoA generated from FAO, we indeed observed a significant elevation of intrahepatic Acetyl-CoA levels from 3 to 12 hours post-PHx (**Fig. 5A**). By using Acetyl-CoA carboxylase inhibitor, firsocostat (FIR), we found that the excessive accumulation of Acetyl-CoA rapidly suppressed the transcription of *Acss2* and *Acly*, while exerting minimal effects on their protein levels (**Fig. S8B and C**). Additionally, our results indicated that both ACLY and ACSS2 are long half-life proteins (**Fig. S8D**). Thus, even with a transient decrease in transcription, they can effectively sustain the conversion and supply of Acetyl-CoA within hepatocytes for up to 12 hours.

**Fig. 5.**
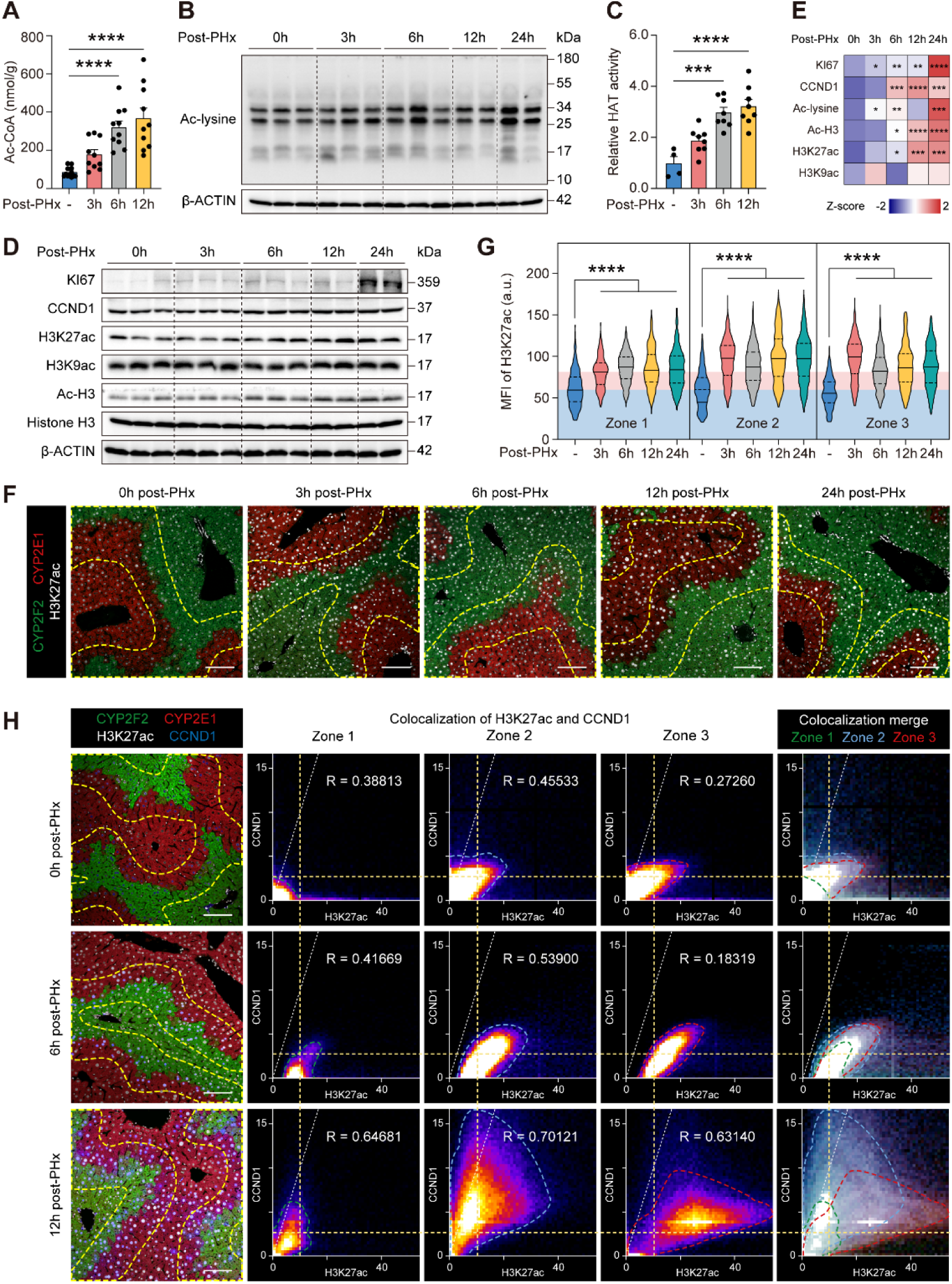
Post-PHx activation of FAO facilitates histone acetylation in the liver. (A) Hepatic Acetyl-CoA levels. (C) HAT activity assay. (B, D-E) Western blotting analysis (B, D) and quantification (E) of Ac-lysine, KI67, CCND1, H3K27ac, H3K9ac, and Ac-H3 in regenerating livers at 0, 3, 6, 12, and 24 hours after liver resection. Histone H3 is used as a loading control for acetylation modifications. β-ACTIN serves as the loading control for other proteins. (F and G) Immunofluorescence staining of liver for CYP2E1, CYP2F2 and H3K27ac. Mean fluorescence intensity (MFI) of H3K27ac in different liver zones is represented in G. Scale bar, 100 μm. CYP2E1^+^ cells in red represent central vein area, while CYP2F2^+^ cells in green represent portal vein area. The yellow dashed lines indicate Zone 2 lobule. Quantifications were performed on at least 5 different areas of the sections in a random way. (H) Staining for CYP2E1, CYP2F2, H3K27ac, and CCND1 on sections from regenerating livers at 0, 6, and 12 hours after PHx (left panel). Scale bar, 100 μm. The scatter plot of colocalization degree between H3K27ac staining and CCND1 staining is calculated by the colocalization finder plug-in in Fiji-ImageJ software (middle panel). R is Pearson’s correlation coefficient and R = 1 indicates complete colocalization (white dotted line). Yellow dotted line represents the leading edge of scatter plot for the control Zone 1 in X- and Y-axis. The green, blue, and red dashed line indicates the leading edge of scatter plot for Zone 1, Zone 2, and Zone 3, respectively. The merged zonal colocalization images are shown in the right panel. \**P*<0.05; \*\**P*<0.01; \*\*\**P*<0.001; \*\*\*\**P*<0.0001, One-way ANOVA. Error bars represent mean ± SEM.

Consistent with the accumulation of Acetyl-CoA, there was a notable upregulation in overall protein acetylation levels (**Fig. 5B**), with particular acetylation of Histones at the 17 kD position, observed from as early as 3 hours post-PHx. This observation is bolstered by the sustained increase in HAT activity throughout the process of liver regeneration (**Fig. 5C**), and the consistent activity of histone deacetylases (**Fig. S8A**). Specifically, the acetylation of H3K27 was significantly more pronounced than that of H3K9, and aligned with the upregulation of proliferation markers (**Fig. 5D and E**). Multiplex immunofluorescence staining further provided deeper insights into the co-localization of H3K27ac with the zonation of proliferative hepatocytes. As shown in **Fig. 5F and G**, the accumulation of H3K27ac was profoundly observed in Zone 2 hepatocytes, and was maintained until 24 hours post-PHx. In contrast, the levels of H3K27ac in Zone 3 peaked at 3 hours post-PHx and gradually declined as liver regeneration progressed. We further observed that the number of H3K27ac and CCND1 double positive hepatocytes was significantly increased in the mid-lobular area (**Fig. 5H**). However, in the periportal and central vein area, although some cells exhibited elevated levels of H3K27ac, there was no increase in CCND1 levels. In the Zone 1 region, the increase in both H3K27ac and CCND1 was not significant. These results further confirm that the spatially specific accumulation of H3K27Ac signature is in accordance with the zonation of liver regeneration.

As expected, by depleting the supply of Acetyl-CoA (**Fig. 6A and Fig. S9A**), ETO treatment and hepatocyte specific CPT1α deletion significantly reduced the acetylation of H3K27, as verified by both WB analysis and multiplex immunofluorescence (**Fig. 6B, C, and J-L** and **Fig. S9B-F**). Similarly, ACLY and ACSS2 inhibitors markedly blunted Acetyl-CoA accumulation and H3K27 acetylation, and subsequently attenuated the initiation of liver regeneration *in vivo*, as characterized by synchronized downregulations in the expression of CCND1 and KI67 (**Fig. S10A-D**). Furthermore, C646, an inhibitor of the paralogue HATs CBP/P300, effectively reduced H3K27 acetylation and eliminated the liver zonation specificity of this acetylation (**Fig. 6F, G, and J-L**). Concurrently, C646 significantly postponed the initiation of liver regeneration, as indicated by the suppressed *Ccnd1* expression and the sharp reduction in KI67- and PCNA-positive hepatocytes in the Zone 2 region (**Fig. 6E, H, and I**). These results suggest that highly activated FAO provides ample substrates for H3K27 acetylation, and are both essential for hepatocytes to re-enter the cell cycle and initiate liver regeneration.

**Fig. 6.**
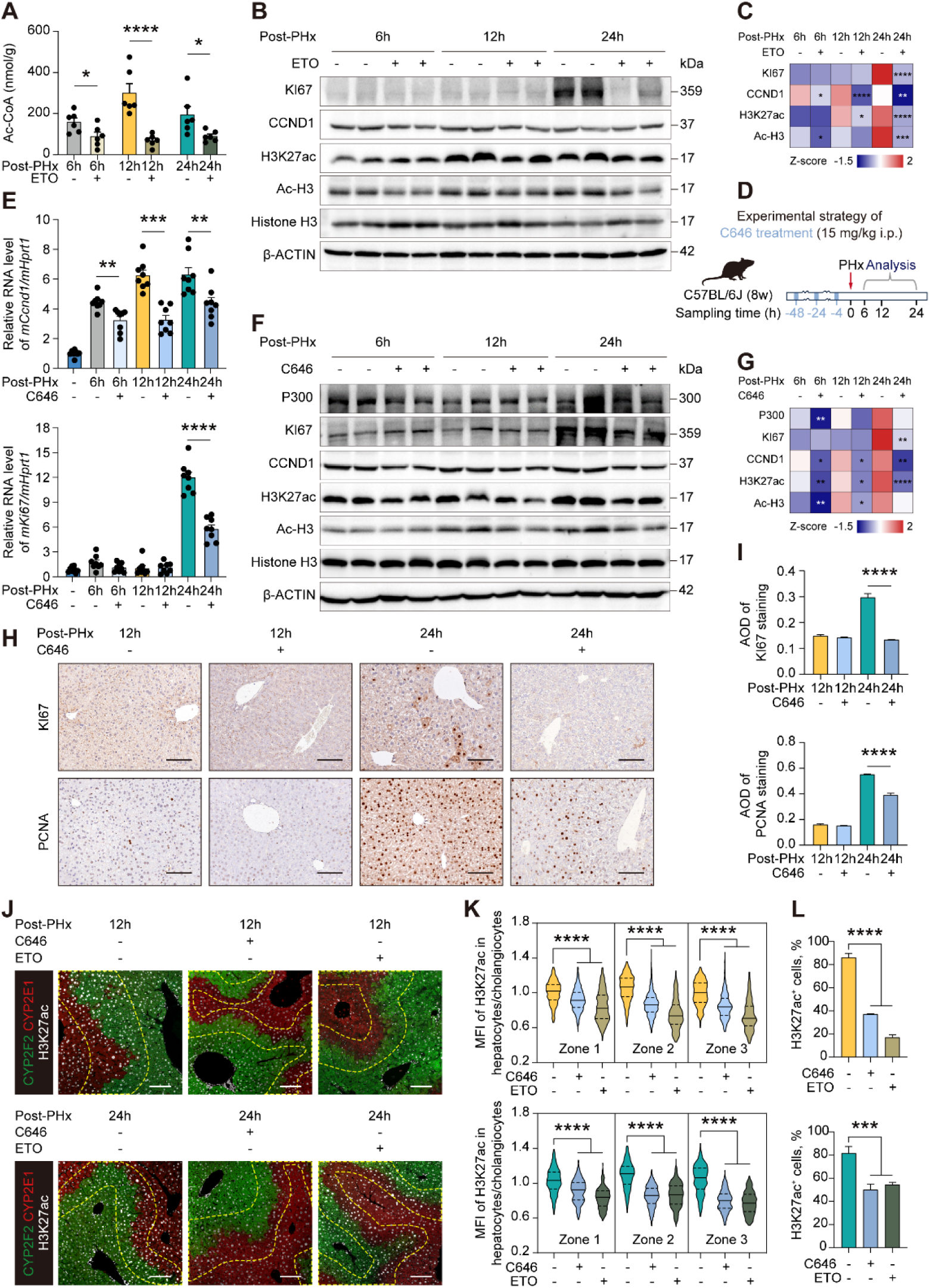
Inhibition of histone acetylation abrogates hepatocyte proliferation. (A) Comparison of hepatic Acetyl-CoA levels with or without ETO treatment. (B and C) Western blotting analysis and quantitation of proliferative hallmarks and histone acetylation modifications in livers sampled at different time points from mice treated with PBS or ETO. H3 was used as a loading control. Histone H3 is used as loading control for acetylation modifications. β-ACTIN serves as loading control for other proteins. (D) Experimental strategy for C646 treatment. (E) qRT-PCR analysis of *Ccnd1* and *Ki67* expression in mouse livers treated with or without C646. (F and G) Western blotting analysis and quantitation of proliferative hallmarks and histone acetylation modifications in livers sampled at different time points from mice treated with PBS or C646. Histone H3 is used as loading control for acetylation modifications. β-ACTIN serves as loading control for other proteins. (H and I) Liver sections were conducted immunohistochemical staining to determine expression of KI67 and PCNA with quantitative analysis of AOD value. Quantifications were performed on at least 5 different areas of the sections in a random way. (J-L) Immunofluorescence staining for CYP2E1, CYP2F2, and H3K27ac on sections from regenerating mice treated with ETO or C646 at 12, and 24 hours after PHx (J). The relative mean fluorescence intensity (MFI) of H3K27ac in hepatocytes compared to cholangiocytes across different liver zonation was represented in K. Proportion of H3K27ac^+^ hepatocytes across sections is in L. Scale bar, 100 μm. CYP2E1^+^ cells in red represent the central vein area, while CYP2F2^+^ cells in green represent the portal vein area. The yellow dashed lines indicate Zone 2 lobule. Quantifications were performed on at least 5 different areas of the sections in a random way. \**P*<0.05; \*\**P*<0.01; \*\*\**P*<0.001; \*\*\*\**P*<0.0001, unpaired Student’s t test or One-way ANOVA. Error bars represent mean ± SEM.

### Enhanced H3K27ac licenses the cell cycle re-entry in hepatocytes

To obtain a landscape view of H3K27 acetylation-mediated epigenetic regulation during the initiation stage of liver regeneration, CUT&Tag assay at 12 hours post-sham or PHx surgery was performed. As shown in **Table S1 and Fig. 7A**, 2,424 peaks with increased H3K27ac signals were identified in the regenerating livers. Notably, we found that H3K27ac signals were enriched around the proximal promoter (<1 kb from the closest TSS) in regenerating liver, characterized by broader distributions (**Fig. S11A and B)**. Next, binding and expression target analysis (BETA) (*26*) was performed to infer target genes, and predicted that differential H3K27ac signatures were correlated with both transcriptional activation (836 genes, termed as up-target) and repression (1036 genes, termed as down-target) (**Table S2 and Fig. 7B**). Pro-proliferative genes, such as *Mcm2*, *Ccne2*, *Cdt1*, *Cks1b*, and *Myc*, were all up-targets (**Fig. 7C**). GOBP pathway analysis further revealed that up-target genes were enriched for mitosis-related terms, such as cell cycle, DNA replication, and cell division, while the down-target genes were involved in lipid metabolism, reflecting a potential feedback mechanism in response to hyperactivation of FAO (**Fig. 7D**). Inspired by the BETA prediction, we identified that 4.2% (101) of genes with increased H3K27ac peaks were transcriptionally activated (**Fig. 7E**). As anticipated, these genes were mainly associated with DNA replication and repair, which are essential for the cell cycle re-entry and mitosis of regenerating hepatocytes, including actin rod assembly, centrosome separation and mitotic sister chromatid cohesion (**Fig. 7F**).

**Fig. 7.**
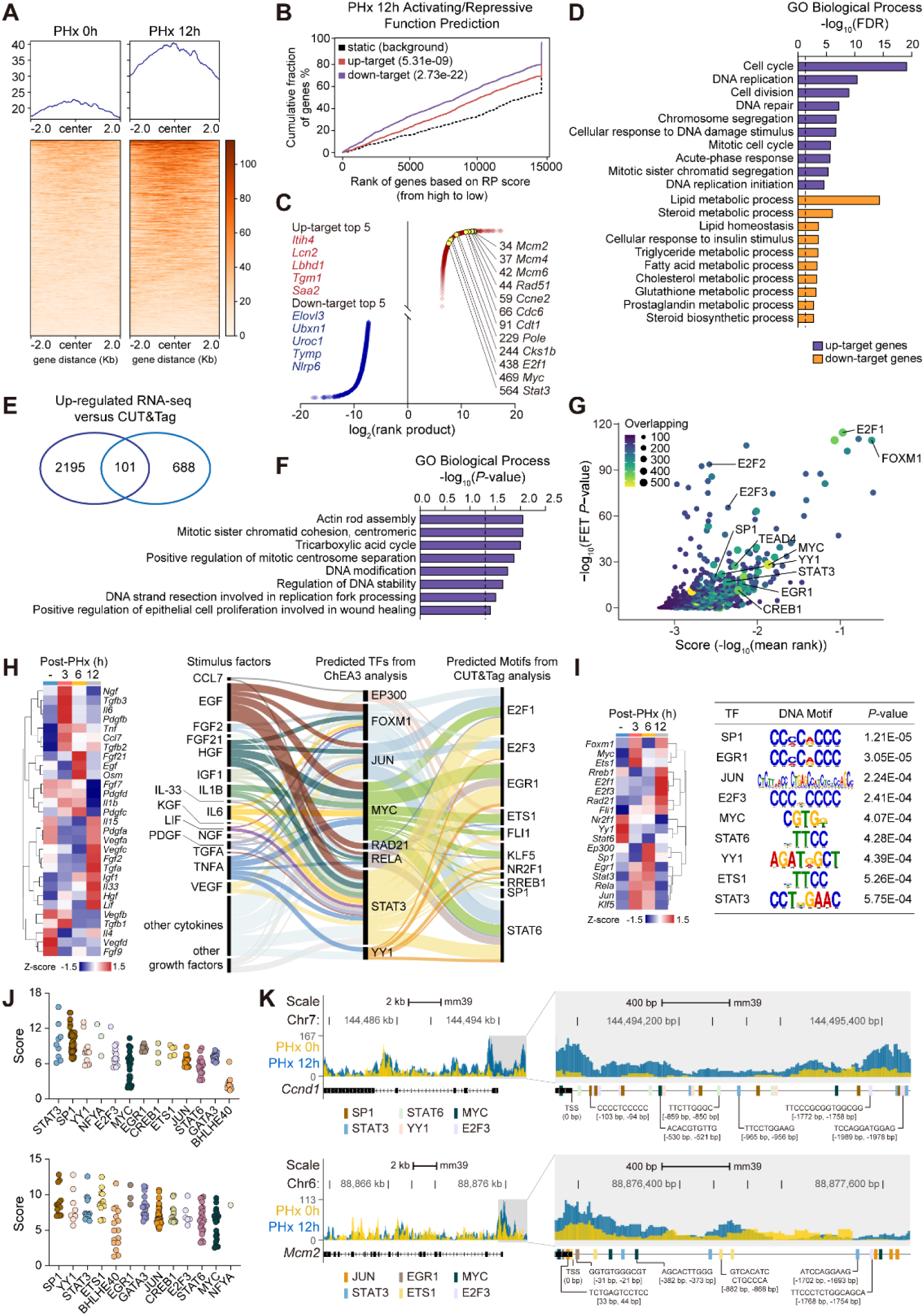
Genome-wide analysis of H3K27ac peaks reveals activation of proliferative pathways. (A) The distribution of peaks according to the CUT &Tag profile of H3K27ac in the group of PHx 0h (left) and PHx 12h (right), n = 2 biological replicates/group. (B) Activating/repressive function prediction by Binding and Expression Target Analysis (BETA). H3K27ac CUT &Tag sites are integrated with RNA-seq gene expression data from livers underwent Sham or PHx surgery. The red and blue lines represent up-target and down-target genes, respectively; dashed lines represent non-DEGs as background. Genes are ranked based on the RP scores of their H3K27ac peaks, and the significance of up- or down-target gene distributions compared to non-DEGs is determined using the Kolmogorov-Smirnov test. RP scores, regulatory potential score. (C) Rank order list of H3K27ac up-target (red dot) and down-target (blue dot) genes in PHx 12h, with proliferative-associated genes highlighted. (D) GOBP pathway analysis for H3K27ac-target genes from **Figure 7C**. (E) Venn diagram showing the overlap between genes with increased expression (789) and genes with elevated H3K27ac deposition (2296) in PHx 12h compared to PHx 0h. (F) Analysis of top GOBP terms from overlapping DEGs. (G) CHIP-X Enrichment Analysis Version 3 (ChEA3) analysis for transcription factor (TF) motif prediction. The genesets used in this analysis are from both up-target genes in **Fig. 7C** and overlapped genes in **Fig. 7E**. (H) Sankey plot summarizing the network of stimulation factors, predicted TFs from ChEA3 analysis, and predicted Motifs from CUT&Tag analysis. The heatmap showing the expression of selected stimulation factors is shown in the left panel. (I) Representative DNA motifs enriched in differentially accessible TFs regions between PHx 12h vs. PHx 0h livers by HOMER motif analysis. Heatmap shows the altered expression of TFs of interests. (J) The JASPAR database (jaspar.genereg.net/) is used to predict the potential TF-binding site on promoter regions of *Ccnd1* and *Mcm2* (threshold > 90%). (K) The distribution of H3K27ac peaks at the genomic loci of *Ccnd1* and *Mcm2* in indicated groups.

ChEA3 analysis based on gene sets identified in both **Fig. 7C and E** was conducted (**Table S3**) and Top 50 predicated transcription factors (TFs) with more than 300 overlapping genes were visualized in **Fig. 7G**. Intriguingly, MEME motif analysis at the peak positions of H3K27ac also predicted that some of these TFs can recognize certain motifs to activate target gene transcription. Therefore, we constructed a stimulation-TFs network to illustrate, under the permissive effects of epigenetic regulation, the signal cascades that promote the transcription initiation of proliferation-related genes. As shown in **Fig. 7H**, the transcription of IL-6, PDGFB and NGF increased after liver resection rapidly, followed by an elevation in EGF, FGF21, and OSM at 6 hours post-PHx, and VEGF and HGF at 12 hours post-PHx. In response to the upregulation of these pro-proliferative factors, MYC and STAT3 emerged as the most extensively connected key nodes among downstream TFs, which could directly recognize H3K27Ac-marked open chromatin regions through corresponding motifs (**Fig. 7I**), thereby promoting transcription. As depicted in **Fig. 7I**, MYC, STAT3, SP1, ETS1, and other TFs were all significantly upregulated from 3 to 12 hours post-PHx, aligning with the overall temporal pattern of transcriptional activation of proliferation-related genes. Two representative cell cycle-related genes, *Ccnd1* and *Mcm2*, were extensively analyzed to validate the aforementioned working model. The JASPAR algorithm predicted that both the promoter regions of *Ccnd1* and *Mcm2* contain abundant and highly credible TF binding elements (**Fig. 7J**). Further visualization revealed that, under PHx induction, H3K27Ac peaks were distributed across the promoter and entire gene body of *Ccnd1*, while for *Mcm2*, they were primarily distributed in the promoter region. These open genomic regions precisely covered multiple repetitive binding elements for TFs such as STAT3, MYC, and E2F3 (**Fig. 7K**). Furthermore, variant enhancer and 797 super-enhancer (SE) loci were identified using the ROSE algorithm (**Fig. S11C**) and were potentially involved in enhancing cell-cycle transition, cytoskeleton remodeling, and mitochondrial respiration (**Fig. S11D and E**). In the predicted SE region for *Ccnd1* and *Cks1b*, significant enrichment of H3K27ac peaks was indeed observed (**Fig. S11F**). Therefore, it can be inferred that the remodeling of H3K27ac profile licenses the free binding of TFs to the promoter region of proliferation-related genes in response to known promoting factors during liver regeneration.

## Discussion

The time-varying characteristics and intrinsic regulatory mechanisms of metabolic and transcriptional bursting during the early phase of liver regeneration following extensive injury and metabolic demands have not been fully understood yet. Our research redefined two distinct periods within the early stage of liver regeneration based on the temporal transcriptome reconstruction that underlies cell cycle re-entry: the preparation phase (3 to 12 hours post-PHx), which have been largely overlooked in previous studies, and the initiation phase (12 to 24 hours post-PHx). Additionally, we profiled the spatiotemporal landscapes of liver metabolic reprogramming, and found a significant shift towards FAO in lipid metabolism during the preparation phase, which coincides with the spatiotemporal pattern of hepatocyte proliferation. In the subsequent initiation phase, there is further rewiring of metabolic replenishment, including increased phospholipid metabolism and glycolysis, to adapt to continuous requirements.

Mounting evidence has shown significant intrahepatic lipid accumulation 24 hours after PHx, leading to debate over whether the imbalance between lipid synthesis and disposal is a consequence of liver repair or a contributing factor to its initiation. Recent studies have shown that active FAO is a common metabolic feature in developing tissues or stem cells with higher regenerative capacity across species (*27*), and is considered as a necessary condition for lymphangiogenesis and heart regeneration (*16, 28*). It is also worth noting that previous studies have indirectly demonstrated that FAO effectively promotes the process of liver regeneration by reshaping the balance between the demand for and supply of fatty acids by inhibiting PTEN or providing ATP and carnitine (*29–31*). Suppressed transient lipid droplet accumulation after PHx resulted in impairment of liver regeneration, with biological implications linked to adipose-derived hormones like adiponectin and leptin, or ATP production (*32, 33*). A most recent publication also suggested that Acetyl-CoA provided by FAO can promote histone acetylation during liver regeneration, but the relevant research was conducted at least 2 days after PHx, which were not be able to provide sufficient evidence in explaining the role of FAO in the initiation of liver regeneration (*34*). However, for the first time, we found that the activation of FAO occurs much earlier than previously reported, representing a salient feature of the preparation phase (3-12 h post-PHx) of liver regeneration. Its significance to host the initiation of liver regeneration is thus unsurprising, although it was long overlooked. Therefore, lipid accumulation or FAB alone may not be the key event that initiates proliferation. Instead, these processes are more likely to provide sufficient fatty acids to fuel FAO, a metabolic adaptation truly associated with the onset of liver regeneration (*35, 36*). This speculation was further supported by the fact that lipid accumulation after ETO treatment or CPT1α deletion hampered hepatocyte proliferation and slowed down liver regeneration, which is consistent with previous studies performed on steatotic liver organoids (*30, 37*). Our data provides the first detailed characterization of FAO among the initial 12 hours post-PHx, and underscores that FAO is indispensable for the initiation of liver regeneration. Hence, facilitating FAO, but not FAB, may represent a novel strategy for large-scale *ex vivo* expansion of liver cells or rescuing regeneration deficient liver in patients. Through CUT&Tag analysis, we have comprehensively examined the mechanisms and target gene sets responsible for H3K27ac remodeling in the initial phases of liver regeneration. Our study also uncovers a previously uncharacterized connection between metabolic reprogramming and epigenetic remodeling, demonstrating how FAO-derived Acetyl-CoA reshapes the epigenetic landscape and guides the re-entry of quiescent hepatocytes into the cell cycle. The rewiring of the epigenome is highly sensitive to the availability of metabolites that serve as substrates for chromatin-modifying enzymes (*38*). During the preparation phase of liver regeneration, hyperactive FAO provides a substantial flux of Acetyl-CoA, which in turn facilitates genome-wide histone acetylation, particularly at H3K27, by upregulating HAT activities. Our results revealed that, similar to the epigenetic modification observed in cancer cell growth, the hijacking of promoters or enhancers by increased deposition of H3K27ac during the initiation phase of liver regeneration supports the expression of proliferation signature genes such as *Myc* and *Ccnd1*, as well as gene sets involved in cytoskeleton reorganization (*39–41*). Our study, for the first time, provides substantial evidence from the perspective of epigenetics to elucidate the causality and underlying mechanisms between FAO and the initiation of liver regeneration, going beyond merely reporting vague correlations as in previous studies. Blocking the FAO process, halting acetyl-CoA generation, or inhibiting HAT activity may alter the direction of metabolic reprogramming during the early stages of liver regeneration, and disrupt its interdependent relationship with epigenetic regulation, leading to slower regeneration and lower survival rate. During a specific stage of stable culture of hepatocyte, providing a supraphysiological supply of intracellular Acetyl-CoA by enhancing FAO or direct supplementation of acetate, and ensure the catalytic efficiency of histone acetylation at the same time, may potentially serve as a promising opportunity to induce reprogramming of hepatocyte into a proliferative state.

The subcellular specific supply of Acetyl-CoA and bulk histone acetylation levels are tightly coordinated through the ACLY-mediated citrate shuttle and ACSS2-depenent acetate conversion (*42*). We further found that the accumulation of Acetyl-CoA inhibited the transcription of these genes. However, both ACLY and ACSS2 are proteins with a long half-life, which allows them to circumvent this negative feedback regulation, ensuring a stable substrate flux for histone acetylation in the nucleus. Although it is challenging to definitively confirm the flow of Acetyl-CoA from FAO to histone acetylation *in vivo* using isotope labeling assay, this aspect can be validated in future studies of *ex vivo* expansion of liver cells. Additionally, the current study found that the acetylation of non-histone proteins was universally upregulated (**Fig. 5B**), suggesting their potential in regulating liver regeneration. Therefore, conducting in-depth research on acetylation proteomics will further enhance our understanding of the relationship between enhanced FAO and the initiation of liver regeneration.

Intriguingly, we also observed a previously uncharacterized self-limiting regulation of metabolic reprogramming, including the dynamic inhibition of FAO and recovery of phospholipid metabolism and FAB occurred 24 hours post-PHx. During the preparation phase of early liver regeneration, when there is no need for phospholipids as structural components of cell membranes, the phospholipid pool is depleted and FAB is suppressed, redirecting the lipid turnover towards FAO. This ensures both efficient energy supply and sufficient substrates for epigenetic modification required for the cell cycle re-entry of hepatocytes. As the liver progresses beyond the initial preparation phase and enters the mitotic process, the proliferating hepatocytes create a huge demand for phospholipids, and the abundance of phospholipids is significantly elevated after a brief downregulation stage. This post-initiation “market-driven” synthesis of phospholipids provides ample substrates for biofilm formation and subsequent organ reconstruction, in accordance with earlier findings of phospholipid metabolism rewired within 48-72 hours after PHx (*14, 43*). Accordingly, reactivated FAB accompanied by downregulated FAO could also jointly ensure the supply of raw materials for phospholipid biosynthesis and accelerate the liver regeneration process (*44, 45*). Here, we also found that this metabolic shift is likely to be influenced by epigenetic modifications, as multiple gene sets involved in fatty acid metabolism, lipid synthesis, and phospholipid metabolism are enriched within the genomic regions exhibiting differential distribution of H3K27ac peaks. Altogether, this epigenetic-related and self-limiting regulation of metabolic reprogramming satisfies the diverse metabolic adaptation needs during different stages of liver regeneration.

In summary, our study uncovers the spatiotemporal landscape of metabolic reprogramming and the genome-wide epigenetic remodeling during the initial 12 hours of liver regeneration, connecting hyperactivated FAO with H3K27ac-dependent transcriptional activation of proliferation-related genes. These findings offer a promising opportunity to manipulate hepatocyte cell fate, potentially driving breakthroughs in regenerative medicine for treating end-stage liver disease.

## Acknowledgements

This work was funded by Excellent Young Scientists Fund, National Natural Science Foundation of China (Grant No. 82322075 to RL) and grants from National Key Research and Development Program on Modernization of Traditional Chinese Medicine (NO. 2022YFC3502104 to RL and XL). XL was supported by the National High-Level Talents Special Support Program. We thank Shanghai Lu Ming Biotech Co., Ltd. (Shanghai, China) for the AFADESI spatial-resolved metabolomics used in this study. We also acknowledge Dr. Ruizhi Li, Yuan Yuan, and Jing Wang for their important suggestions and valuable technical support.

## Availability of Data and Materials

All data associated with this study are included within the main text or the **Supplementary information**. Raw RNA-seq data, Untargeted Metabolomics data, Spatial Metabolomics data, and CUT&Tag data generated in the present study have been deposited at NCBI GEO databank and other public databases and will be publicly available as of the date of publication. Other data reported in this paper will be shared by the corresponding author upon request.

## Authors contributions

Study concept and design: RL; acquisition of data: QZ, XL, YC, FL, XX, SL, RS, GF, JW, JQ; analysis and interpretation of data: QZ, XL, ZX; drafting of the manuscript: QZ, RL; critical revision of the manuscript for important intellectual content: XL, RL; statistical analysis: QZ; obtained funding: RL, XL; administrative, technical, or material support: ZX, XL, RL; study supervision: RL. All authors approved the final version of the manuscript.

## Abbreviations

2-DG: 2-deoxy-D-glucose
Acars: acylcarnitines
ACC: acetyl-CoA carboxylase
ACLY: ATP citrate lyase
ACOX1: acyl-CoA oxidase 1
ACSS2: acyl-CoA synthetase short chain family member 2
AFADESI-MSI: airflow-assisted desorption electrospray ionization mass spectrometry imaging
CPT1A: carnitine palmitoyltransferase 1A
DEGs: differentially expressed genes
ETO: etomoxir
FAB: fatty acid biosynthesis
FAO: fatty acid oxidation
FIR: firsocostat
GOBP: gene ontology biological processes
H&E: hematoxylin and eosin
HAT: histone acetyltransferase
HCA: hierarchical clustering analysis
HK2: hexokinase 2
LC-MS/MS: liquid chromatography coupled to tandem mass spectrometry
LDHA: lactate dehydrogenase A
OXA: sodium oxamate
PC: phosphatidylcholine
PE: phosphatidylethanolamine
PHx: partial hepatectomy
SE: super-enhancer
TDYA: 10,12-tricosadiynoic acid
TFs: transcription factors
TG: triglycerides

