## Supplementary materials for "Temporal-spatially metabolic reprogramming rewires the H3K27ac landscape to enable the initiation of liver regeneration"

### Supplementary methods

#### Cell line and treatments

Human hepatoblastoma (HepG2) cell line was plated and cultured in DMEM medium supplemented with 10% fetal bovine serum, and 1% (v/v) penicillin/streptomycin at 37 °C and 5% $CO_2$. One hour before the cell treatment, the culture medium was changed to a medium with 1% fetal bovine serum. For time-course treatment, cells were treated with Firsocostat (500 mM) and harvested for RNA or protein extraction at different time points. For the half-life assay, cells were treated with Cycloheximide (100 mM) and collected at the indicated time points and prepared for western blotting analysis.

#### Acetyl-CoA assay

Acetyl-CoA was measured using an Acetyl-CoA assay kit (BC0980, Solarbio) in accordance with the manufacturer's instructions. The snap frozen liver samples were homogenized in lysis buffer of the kit on ice. The supernatant was used to determine Acetyl-CoA concentration in duplicate according to manufacturer’s instructions, and the OD value for spectrophotometric measurements was read at 340 nm at 20 sec and 80 sec after adding the working solution to the wells, respectively. Acetyl-CoA content was calculated as the difference between OD values at different times divided by the sample weight.

#### HAT and HDAC activity assay

The liver protein was extracted and quantitated using the BCA protein assay kit (BN27109, Biorigin). The HAT activity assay was carried out by utilizing HAT activity assay kit (ab65352, Abcam), and the HDAC activity was determined using HDAC assay kit (56210, ACTIVE MOTIF). The absorbance was read at 440 nm and 405 nm to measure the OD value, respectively. Both HAT and HDAC activity were shown as the percentage of control group.

**Histology and immunohistochemistry**

4% Paraformaldehyde solution was used to fix liver tissues for 1 week. Paraffin-embedded sections (4 μm thick) were deparaffinized, rehydrated, and antigen-retrieved. For H&E staining, slides were stained with Mayer's hematoxylin and 0.1% sodium bicarbonate, and then counterstained with Eosin Y solution. For immunohistochemistry, slides were washed, blocked in 10% goat serum, and incubated with the corresponding primary antibodies at 4 °C overnight. The chromogenic reaction was performed using a DAB solution in the immunohistochemical staining process. Slides were scanned using the Leica Aperio VERSA Digital Pathology Scanner (Aperio Technologies Inc.). Multiplex immunofluorescence was performed with sequential incubation of different primary antibodies, followed by washes and incubation with the secondary antibody and TSA reagents with different fluoresceins. With the accumulation of fluorescent signals, the previous round of antibodies was stripped, followed by the incubation of the next round of antibodies. Nuclei were stained with DAPI after all the antigens were labeled. Images were acquired using a confocal microscope (Lecia TCS SP8, Lecia Stellaris). The average optical density (AOD) of immunohistochemical and mean fluorescence intensity (MFI) of multiplex immunohistochemistry was analyzed with ImageJ software (Wayne Rasband National Institutes of Health, USA). The antibodies used in the immunohistochemistry and multiplex immunohistochemistry included anti-PCNA (1:5000, Cell Signaling Technology, 2586s), anti-KI67 (1:10000, proteintech, 27309-1-AP), anti-CCND1 (1:200, Abcam, ab134175), anti-H3K27ac (1:100, Cell Signaling Technology, 4353S), anti-CYP2E1 (1:100, proteintech, 19937-1-AP), anti-CYP2F2 (1:60, SantaCruz, sc-374540).

**Western blotting analysis**

Protein extraction from liver tissues and HepG2 cells was conducted using RIPA Lysis Buffer and was quantified by BCA protein assay kit (Biorigin (Beijing) Inc., Beijing, China). Protein lysates were then analyzed by SDS-PAGE and transferred to the PVDF membrane for immunoblotting. Proteins of interest were detected by incubating membranes at 4 °C overnight in blocking buffer with the indicated primary antibody. After washing three times with TBST, membranes were probed with appropriate secondary antibodies for 1 hour, washed again, detected by using Super ECL Plus Western Blotting System Kit (Biorigin (Beijing) Inc., Beijing, China). Blotting signaling was visualized with Tanon-5200 Chemiluminescent Imaging System (Tanon, Shanghai, China). Densitometry results of western blotting were quantified using ImageJ software. Following antibodies were used: anti-KI67 (1:5000, proteintech, 27309-1-AP), anti-CCND1 (1:5000, proteintech, 60186-1-lg), anti-ACSS2 (1:200, Santa Cruz, sc-398559), anti-ACLY (1:200, Santa Cruz, sc-517267), anti-H3K27ac (1:1000, Cell Signaling Technology, 4353S), anti-Histone-H3 (1:1000, proteintech, 17168-1-AP), anti-Ac-H3 (1:400, Santa Cruz, sc-56616), anti-Ac-lysine (1:150, Santa Cruz, sc-32268), and anti-β-Actin (1:5000, proteintech, 66009-1-lg).

**RNA sequencing**

Total RNA was extracted from mice livers using TRIzol reagent. Approximately 1 μg of total RNA extracted from mouse livers was used for RNA-seq library preparation following the Illumina TruSeq stranded mRNA sample preparation guide (Illumina, San Diego, CA). Briefly, poly-A-containing mRNA molecules were first purified using poly-T oligo-attached magnetic beads. The purified mRNA was then fragmented and converted into first-strand cDNA using reverse transcriptase and random primers. This was followed by second-strand cDNA synthesis using RNase H, DNA polymerase I, and dNTP. After adapter ligation, products were then purified and enriched by PCR amplification to generate the final RNA-seq library.

The RNA-seq libraries were subjected to quantification process, pooled for cBot amplification, and subsequently sequenced on an Illumina NovaSeq 6000 platform with 150 bp paired-end reads. FastQ files were generated containing nucleotide data and quality scores for each position. Sequencing reads were mapped to the *Mus musculus* reference genome GRCm38 using Hisat2 software (version 2.0.5). Clean reads were quantified based on the length and reads count mapped to this gene, and their FPKM were calculated using featureCounts (version 1.5.0-p3). Aligned BAM files were sorted by SAMtools. DESeq2 R packages (version 1.20.0) was used for read counting and differential expression analysis. False discovery rate (FDR) correction was performed using Benjamini and Hochberg's approach to obtain a P correction. FDR ≤ 0.05 and $|log_2FC| \geq 1$ were set as the threshold for significantly differential expression. GO and KEGG enrichment analysis were performed on DEGs. For functional profiling of DEGs, GSEA analysis was performed. The raw RNA-seq data were uploaded to the GEO database with accession number GSE255995 and GSE256002.

**Untargeted Metabolomics**

Tissues (~100 mg) were individually grounded with liquid nitrogen and the homogenate was resuspended with prechilled 80% methanol and 0.1% formic acid by well vortex. The samples were incubated on ice for 5 min and then were centrifuged at 15,000 g, 4 °C for 20 min. Some of supernatant was diluted to final concentration containing 53% methanol by LC-MS grade water. The samples were subsequently transferred to a fresh Eppendorf tube and then were centrifuged at 15000 g, 4 °C for 20 min. Finally, the supernatant was injected into the LC-MS/MS system analysis [1]. LC-MS/MS analyses were performed using a Vanquish UHPLC system (Thermo Fisher, Germany) coupled with an Orbitrap Q Exactive$^{TM}$ HF mass spectrometer (Thermo Fisher, Germany). Samples were injected onto a Hypesil GOLD column (100×2.1 mm, 1.9 μm) using a 12-min linear gradient at a flow rate of 0.2 mL/min. The eluents for the positive polarity mode were eluent A (0.1% FA in Water) and eluent B (Methanol). The eluents for the negative polarity mode were eluent A (5 mM ammonium acetate, pH 9.0) and eluent B (Methanol). The solvent gradient was set as follows: 2% B, 1.5 min; 2-85% B, 3.0 min; 85-100% B, 10.0 min; 100-2% B, 10.1 min; 2% B, 12 min. Q Exactive$^{TM}$ HF mass spectrometer was operated in positive/negative polarity mode with spray voltage of 3.5 kV, capillary temperature of 320 °C, sheath gas flow rate of 35 psi and aux gas flow rate of 10 L/min.

Samples were analyzed in a randomized fashion and QC samples were additionally measured in confirmation mode to obtain additional MS/MS spectra for identification. The raw data files were processed using the Compound Discoverer 3.1 (Thermo Fisher) to perform peak alignment, peak picking, and quantitation for each metabolite. The main parameters were set as follows: retention time tolerance, 0.2 minutes; actual mass tolerance, 5 ppm; signal intensity tolerance, 30%; signal/noise ratio, 3; and minimum intensity, 100,000. After that, peak intensities were normalized to the total spectral intensity. The normalized data was used to predict the molecular formula based on additive ions, molecular ion peaks and fragment ions. And then peaks were matched with the mzCloud (https://www.mzcloud.org/), mzVault and MassList database to obtain the accurate qualitative and relative quantitative results.

These peaks were then identified and validated by aligning the molecular mass data (m/z) using the online KEGG (http://www.genome.jp/kegg/), HMDB (https://hmdb.ca/) and LIPID Maps (https://www.lipidmaps.org/) database. We applied t-test analysis to calculate the statistical significance. The metabolites with VIP > 1 and $P < 0.05$ and FC ≥2 or FC ≤ 0.5 were considered to be differential metabolites. The functions of these metabolites and metabolic pathways were studied using the KEGG database. The metabolic pathways enrichment of differential metabolites was performed, when ratio was satisfied by x/n > y/N, the metabolic pathway was considered as enrichment, when P of metabolic pathway < 0.05, the metabolic pathway was considered as statistically significant enrichment. The processed untargeted metabolome data were uploaded to the OMIX database (China National Center for Bioinformation/Beijing Institute of Genomics, Chinese Academy of Sciences), under accession number OMIX005833 and OMIX005834.

**Airflow-assisted desorption electrospray ionization mass spectrometry imaging analysis**

The embedded liver samples were cut into consecutive sagittal slices 10 µm about 10 slices using a cryostat microtome (Leica CM 1950, Leica Microsystem, Germany) and were then thaw-mounted onto positive charge desorption plate (Thermo Scientific, U.S.A). They were desiccated at -20 °C for 1 h and then at room temperature for 2 h before MSI analysis. Meanwhile, an adjacent slice was left for H&E staining to confirm the liver zonation. The analysis was performed as previously reported [2]. In brief, this experiment was carried out with an AFADESI-MSI platform (Beijing Victor Technology Co., LTD,

Beijing, China) in tandem with a Q-Orbitrap mass spectrometer (Q Exactive, Thermo Scientific, U.S.A.). Here, the solvent formula was acetonitrile (ACN) /H2O (8:2) at negative mode and ACN/H2O (8:2, 0.1%FA) at positive mode and the solvent flow rate was 5 μL/min, the transporting gas flow rate was 45 L/min, the spray voltage was set at 7 kV, and the distance between the sample surface and the sprayer was 3 mm as was the distance from the sprayer to the ion transporting tube. The MS resolution was set at 70,000, the mass range was 70-1000 Da, the automated gain control target was 2E6, the maximum injection time was set to 200 ms, the S-lens voltage was 55 V, and the capillary temperature was 350 °C. The MSI experiment was carried out with a constant rate of 0.2 mm/s continuously scanning the surface of the sample section in the X direction and a 40 μm vertical step in the Y direction.

The collected .raw files were converted into .imML format using imzMLConverter and then imported into MSiReader for ion image reconstructions after background subtraction using the Cardinal software package. All MS images were normalized using total ion count normalization in each pixel. Region-specific MS profiles were precisely extracted by matching high-spatial resolution H&E images. The discriminating endogenous molecules of different tissue microregions were screened by a supervised statistical analytical method: orthogonal partial least squares discrimination analysis (OPLS-DA). VIP values obtained from the OPLS-DA model were used to rank the overall contribution of each variable to group discrimination. The VIP value reflects the importance degree on the classification of sample categories with respect to the first two principal components of the OPLS-DA model, which indicates that this variable has a significant effect if the VIP is greater than 1. A two-tailed Student's T-test was further used to verify whether the metabolites of difference between groups were significant. Differential metabolites were selected with VIP values greater than 1.0 and $P < 0.05$. Additionally, for the special data structure obtained from the MSI analysis, we also performed T-distributed stochastic neighbor embedding on the MS data in each pixel for dimensionality reduction, respectively. The Spatial shrunken centroids clustering was applied for MSI data clustering to separate the sample based on the differences of abundance of ions in each pixel. The ions detected by AFADESI were annotated by the pySM pipeline and an in-house SmetDB database (Lumingbio, Shanghai, China). AFADESI-MSI data have been uploaded to METASPACE database (https://metaspace2020.eu/) under the project name "Liu et al. (2024) liver regeneration post-70%PHx".

**CUT&Tag analysis**

CUT&Tag assay was performed using Hyperactive™ In-Situ ChIP Library Prep Kit for Illumina (TD901-TD902, Vazyme Biotech, China) according to manufacturer's instruction [3]. Briefly, prepared concanavalin A-coated magnetic beads (ConA beads) were added to resuspended cells and incubated at room temperature to bound cells. Non-ionic detergent Digitonin was used to permeate cell membrane. Then, primary H3K27ac antibody (ab177178,

Abcam), secondary antibody and the Hyperactive pA-Tn5 Transposase were incubated with the cells that were bounded by ConA beads in order. Therefore, the Hyperactive pA-Tn5 Transposase can exactly cut off the DNA fragments that were bound with target protein. In addition, the cut DNA fragments can be ligated with P5 and P7 adaptors by Tn5 transposase and the libraries were amplified by PCR with the P5 and P7 primers. The purified PCR products were evaluated using the Agilent 2100 Bioanalyzer (Agilent Technologies, Santa Clara, CA, USA). Finally, these libraries were sequenced on the Illumina NovaSeq6000 platform and 150bp paired-end reads were generated for the following analysis.

The raw sequence data were firstly quality trimmed by fastq software to obtain the clean reads. Then the clean reads were aligned to the *Mus musculus* genome GRCm39 using Bowtie2 and subsequently analyzed by the SEACR software based the 'stringent' parameter to detect genomic regions enriched for multiple overlapping DNA fragments (peaks) that we considered to be putative binding sites. Visualization of peak distribution along genomic regions of interested genes was performed with IGV. Peaks were then annotated using chipseeker software to obtain the genes and gene annotations about peaks. Significant Motif of peaks were analyzed by MEME and DREME software and aligned to the motif database. Different peaks between case (12 hours post-PHx) and control groups were analyzed using Diffbind software. Firstly, the overlapping and merging of peaks were processed and the reads counts of peaks was calculated. Then, the statistically significant differential binding sites were identified based on binding affinity with the threshold of $P < 0.05$ and $|\log_2 FC| > 1$. CUT&Tag sequencing data are available under the GEO accession number GSE255292.

**Quantification and visualization of multi-omics data**

For multi-omics sequencing analysis, a minimum of three biological replicates were analyzed, with the exception of only two biological replicates for CUT&Tag. Sequencing data analysis was performed using R package software (version 4.3.2). The PCA and Volcano plot were generated by R package ggplot2 (3.4.4) and ggrepel (0.9.4). Cluster membership and hierarchical heatmap-based visualization for the temporal dynamics of gene expression were plotted using the R packages: ComplexHeatmap (2.18.0) and Mfuzz (2.62.0). Pearson correlation coefficient values were computed using the corr.test function in package psych (2.3.9), and were visualized through reshape2 (1.4.4) and pheatmap (1.0.12) in R software.

### Supplementary figures

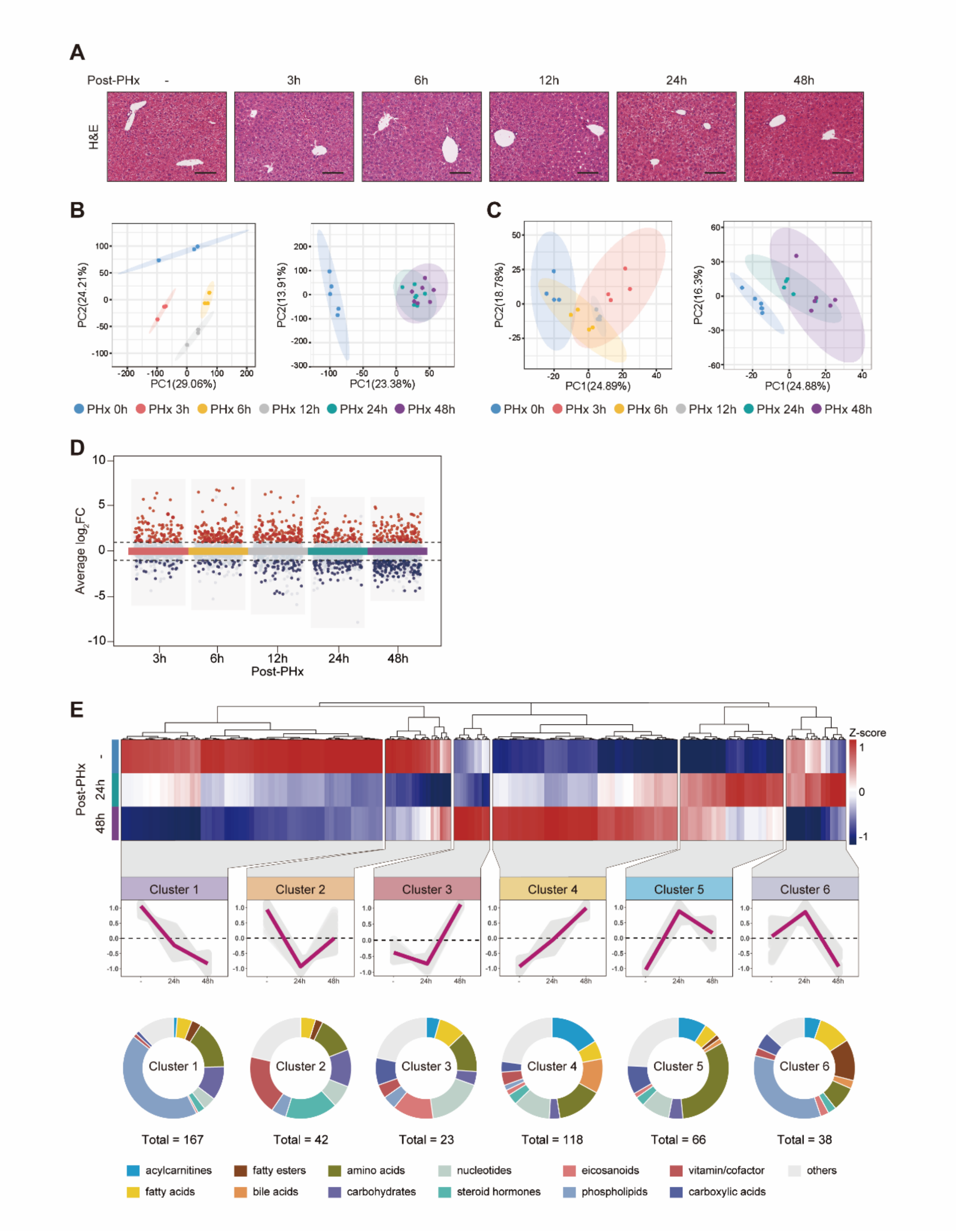

**Fig. S1. Overview of transcriptome and untargeted metabolome analysis for regenerating liver after 70% PHx.**

(A) Representative images of H&E staining for regenerating liver at different times. Scale bars, 100 μm.

(B) Transcriptome-based principal component analysis (PCA) revealed the distinct developmental trajectories in the priming phase (left) and regenerating phase (right) of early liver regeneration.

(C) PCA projection of metabolomic profiles in the early liver regeneration.

(D) Metabolomics volcano plots illustrating the temporal changes in metabolite content in terms of statistical significance versus magnitude of change. Red symbols classify the upregulated metabolites, while the blue represents downregulated ones, according to the criteria: $|\log_2 FC| > 1$ and $FDR < 0.05$. Dashed lines represent the average $\log_2 FC$ value ± 1, respectively.

(E) Heatmap shows the relative abundance of metabolites involved in 24-48 hours post-PHx. 6 clusters were highlighted, and the trend lines indicating the change of intrahepatic levels of these metabolites were shown. Metabolites class compositions in each cluster were subcategorized and represent in pie-chart.

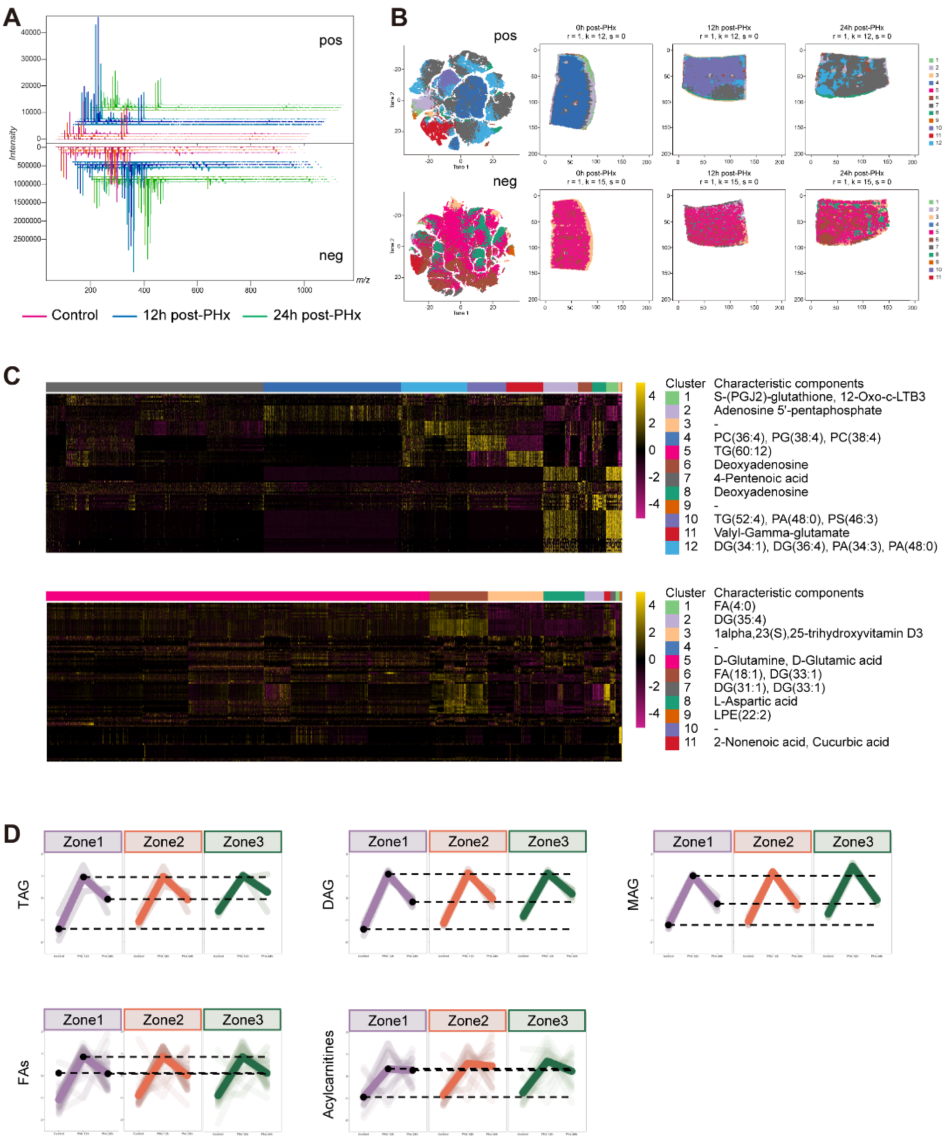
A
pos
neg
Intensity
m/z
Control
12h post-PHx
24h post-PHx
B
pos
neg
0h post-PHx
12h post-PHx
24h post-PHx
C
Cluster Characteristic components
1 S-(PGJ2)-glutathione, 12-Oxo-c-LTB3
2 Adenosine 5'-pentaphosphate
3 -
4 PC(36:4), PG(38:4), PC(38:4)
5 TG(60:12)
6 Deoxyadenosine
7 4-Pentenoic acid
8 Deoxyadenosine
9 -
10 TG(52:4), PA(48:0), PS(46:3)
11 Valyl-Gamma-glutamate
12 DG(34:1), DG(36:4), PA(34:3), PA(48:0)
Cluster Characteristic components
1 FA(4:0)
2 DG(35:4)
3 1alpha,23(S),25-trihydroxyvitamin D3
4 -
5 D-Glutamine, D-Glutamic acid
6 FA(18:1), DG(33:1)
7 DG(31:1), DG(33:1)
8 L-Aspartic acid
9 LPE(22:2)
10 -
11 2-Nonenoic acid, Cucurbic acid
D
Zone1
Zone2
Zone3
TAG
DAG
MAG
FAs
Acylcarnitines

**Fig. S2. AFADESI-MSI analysis for regenerating liver after 70% PHx.**

(A) Extracted time-specific metabolite MS profile.

(B and C) TSNE analysis, cluster heatmap analysis and characteristic components of specifically enriched metabolites in different liver zonation micro-regions.

(D) Statistics highlighting the spatial distribution of metabolites in **Figure 3C**.

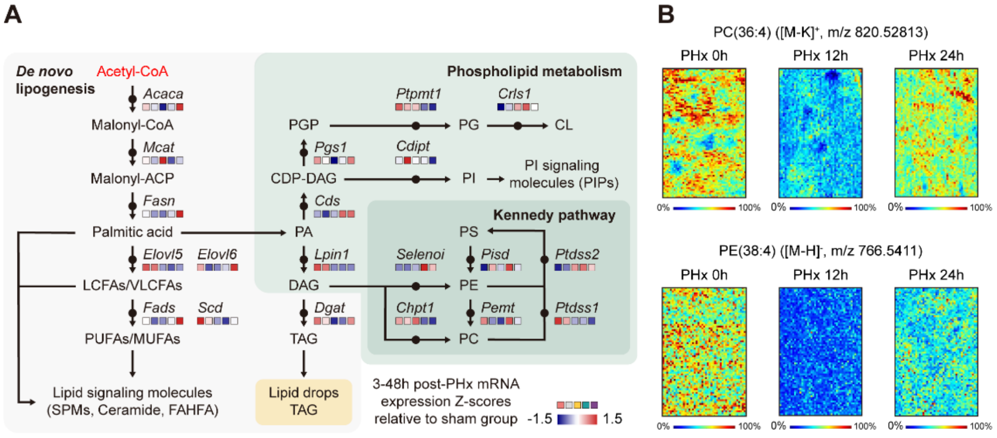
A
De novo lipogenesis
Acetyl-CoA
Acaca
Malonyl-CoA
Mcat
Malonyl-ACP
Fasn
Palmitic acid
Elovl5
Elovl6
LCFAs/VLCFAs
Fads
Scd
PUFAs/MUFAs
Lipid signaling molecules
(SPMs, Ceramide, FAHFA)
Phospholipid metabolism
Ptpmt1
Crls1
PGP
PG
CL
Pgs1
Cdipt
CDP-DAG
PI
PI signaling
molecules (PIPs)
Cds
PA
Lpin1
DAG
Dgat
TAG
Lipid drops
TAG
Kennedy pathway
PS
Selenoi
Pisd
Ptdss2
PE
Chpt1
Pemt
Ptdss1
PC
3-48h post-PHx mRNA
expression Z-scores
relative to sham group
-1.5
1.5
B
PC(36:4) ([M-K]+, m/z 820.52813)
PHx 0h
PHx 12h
PHx 24h
PE(38:4) ([M-H]-, m/z 766.5411)
0%
100%

**Fig. S3. *De novo* lipogenesis and phospholipid metabolism in the initiation phase of liver regeneration.**

(A) Fatty acids are synthesized by *de novo* lipogenesis (*Acaca* and *Fasn*), and subsequently elongated (ELOVL family) and desaturated (*Fads* and *Scd*). Fatty acids and glycerol are assembled into triglycerides (LPIN and DGAT family). The phospholipid-synthetic pathways are branched into several different directions, leading to phosphatidylglycerol (PG), cardiolipin (CL), phosphatidylinositol (PI), phosphatidylcholine (PC), phosphatidylethanolamine (PE), and phosphatidylserine (PS). These pathways are interconnected and constantly in flux. FPKM at different time points from 3 to 48 hours were normalized relative to FPKM of the sham group. Z-scores were visualized with color ranging from red (high expression relative to sham) over white (intermediate expression relative to sham) to blue (low expression relative to sham).

(B) MSI images of representative PC and PE in liver tissues (intensity in color scale is relative value).

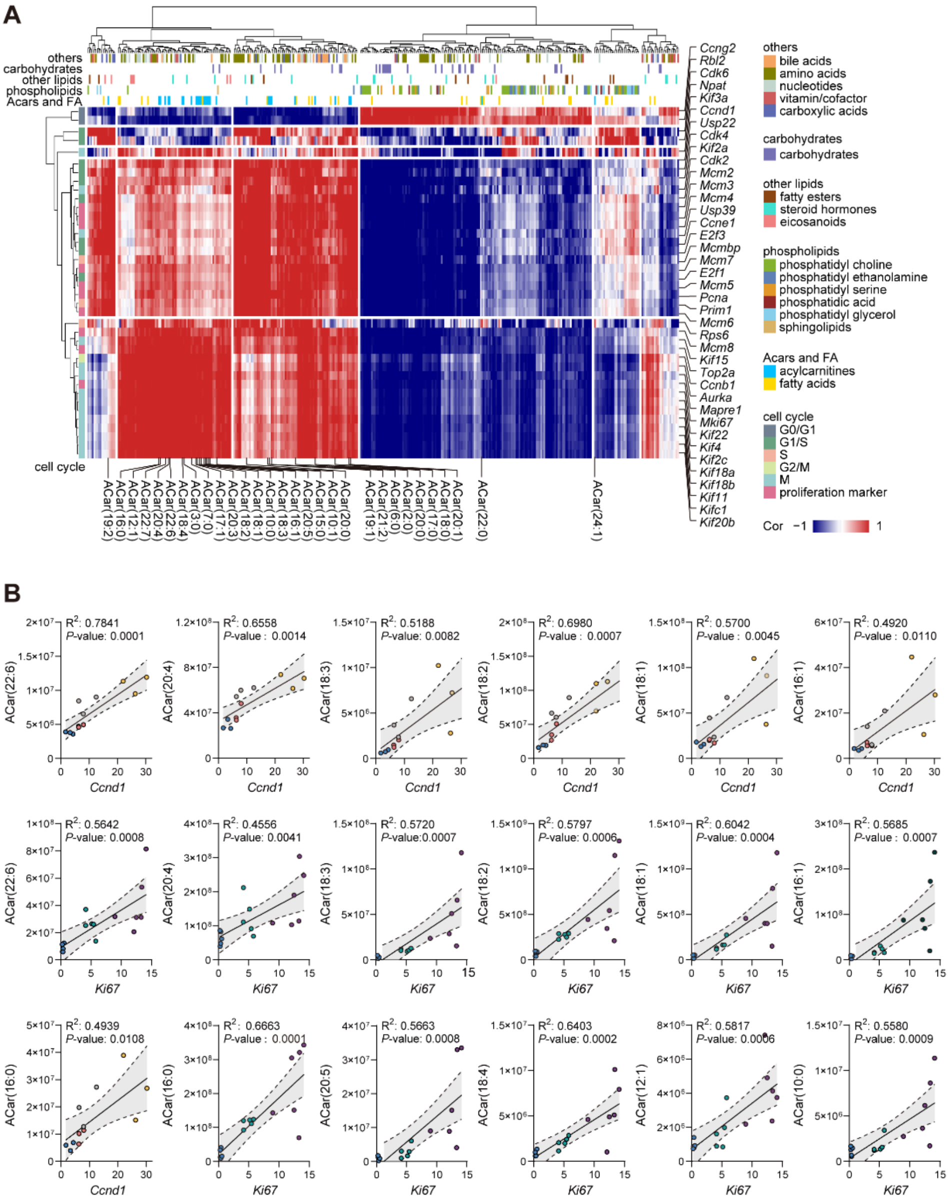

**Fig. S4. The correlation analysis between rewired lipid metabolism and hallmarks of proliferation in the initiation phase of liver regeneration.**

(A) Correlation analysis of the association between the proliferating genes and differential metabolites at 24 to 48 hours post-PHx. The Pearson's correlation (R) values were visualized in heatmap. Each kind of metabolites analyzed are annotated above the heatmap with details shown in the legend to the right. The color code representing each cell cycle phase is shown on the lower right legend of the heatmap.

(B) Scatter plot and simple linear regression demonstrating association between Acars levels and *Ccnd1* or *Ki67* expression. The correlation coefficient ($R^2$) and *P* from simple linear regression are shown.

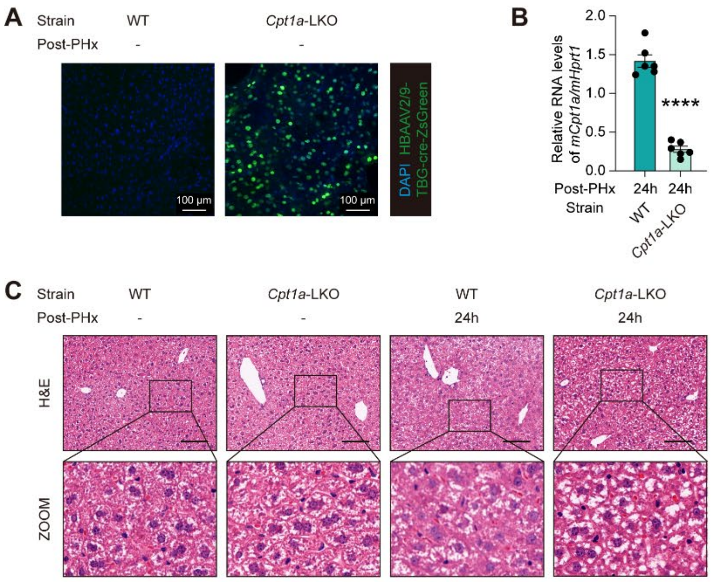
A
Strain
WT
Cpt1a-LKO
Post-PHx
-
-
100 μm
100 μm
DAPI HBAAV2/9-TBG-cre-ZsGreen
B
Relative RNA levels of mCpt1a/mHprt1
2.0
1.5
1.0
0.5
0.0
****
Post-PHx
24h
24h
Strain
WT
Cpt1a-LKO
C
Strain
WT
Cpt1a-LKO
WT
Cpt1a-LKO
Post-PHx
-
-
24h
24h
H&E
ZOOM

**Fig. S5. The construction and verification of *Cpt1a*-LKO mouse model.**

(A) Representative immunofluorescence images of ZsGreen expression in the liver after tail vein injection of HBAAV2/9-TBG-cre-ZsGreen viruses. Scale bar, 100 μm.

(B) The transcription levels of *Cpt1a* in livers of WT and *Cpt1a*-LKO mice at 24 hours post-PHx were determined by qRT-PCR assay. Data are represented as mean ± SEM of 6 replicates.

(C) H&E staining of liver tissues. Scale bars, 100 μm. The lower panel shows a zoom-in view of the area in a black box in the upper panel.

$^{*}P<0.05$; $^{**}P<0.01$; $^{***}P<0.001$; $^{****}P<0.0001$, unpaired Student's t test. Error bars represent mean ± SEM.

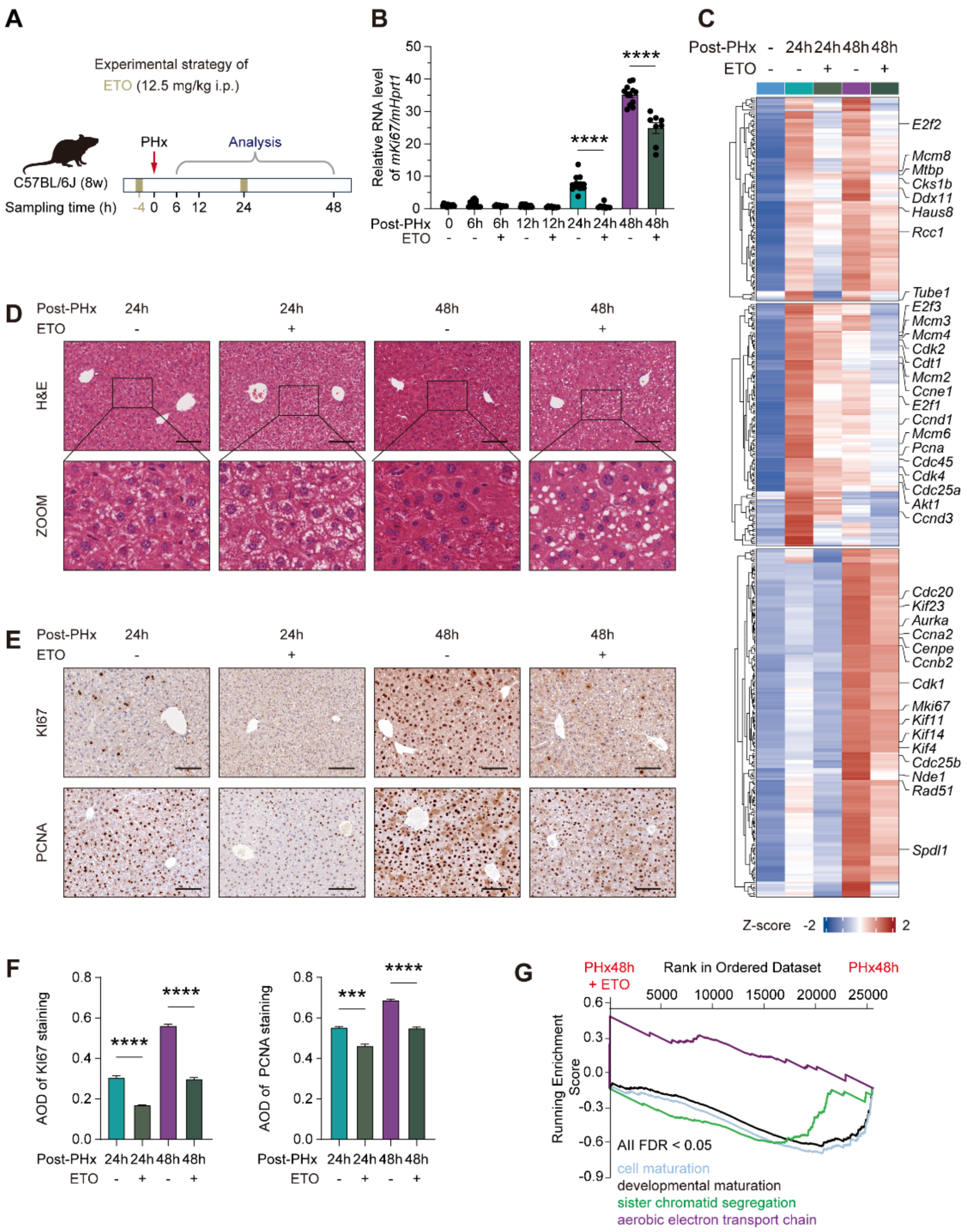
A
Experimental strategy of
ETO (12.5 mg/kg i.p.)
PHx
Analysis
C57BL/6J (8w)
Sampling time (h) -4 0 6 12 24 48
B
Relative RNA level of mKi67/mHprt1
Post-PHx 0 6h 6h 12h 12h 24h 24h 48h 48h
ETO - - + - + - + - +
C
Post-PHx - 24h 24h 48h 48h
ETO - - + - +
E2f2
Mcm8
Mtbp
Cks1b
Ddx11
Haus8
Rcc1
Tube1
E2f3
Mcm3
Mcm4
Cdk2
Cdt1
Mcm2
Ccne1
E2f1
Ccnd1
Mcm6
Pcna
Cdc45
Cdk4
Cdc25a
Akt1
Ccnd3
Cdc20
Kif23
Aurka
Ccna2
Cenpe
Ccnb2
Cdk1
Mki67
Kif11
Kif14
Kif4
Cdc25b
Nde1
Rad51
Spdl1
Z-score -2 2
D
Post-PHx 24h 24h 48h 48h
ETO - + - +
H&E
ZOOM
E
Post-PHx 24h 24h 48h 48h
ETO - + - +
KI67
PCNA
F
AOD of KI67 staining
AOD of PCNA staining
Post-PHx 24h 24h 48h 48h
ETO - + - +
G
PHx48h + ETO
Rank in Ordered Dataset
PHx48h
Running Enrichment Score
All FDR < 0.05
cell maturation
developmental maturation
sister chromatid segregation
aerobic electron transport chain

**Fig. S6. The CPT1A inhibitor ETO inhibits cell cycle progression and cell proliferation after PHx.**

(A) Animal experimental strategy of inhibitor treatment.

(B) The transcription levels of *Ki67* were determined by qRT-PCR assay. Data are represented as mean ± SEM of 8 or 12 replicates.

(C) Heatmap of mRNA expression changes after ETO treatment relative to PBS-treated mice according the color scale shown.

(D) H&E staining of liver tissues. Scale bars, 100 μm. The lower panel shows a zoom-in view of the area in a black box in the upper panel.

(E and F) Immunohistochemical staining determines the expression of KI67 and PCNA, with quantitative analysis of AOD value. Quantifications were performed on at least 5 different areas of the sections in a random way.

(G) GSEA enrichment plots showing a significant correlation (determined by FDR < 0.05) of down-regulated genes in the ETO-treated livers with hallmark data associated with proliferation pathways. FDR, false discovery rate.

$^{*}P<0.05$; $^{**}P<0.01$; $^{***}P<0.001$; $^{****}P<0.0001$, unpaired Student's t test. Error bars represent mean ± SEM.

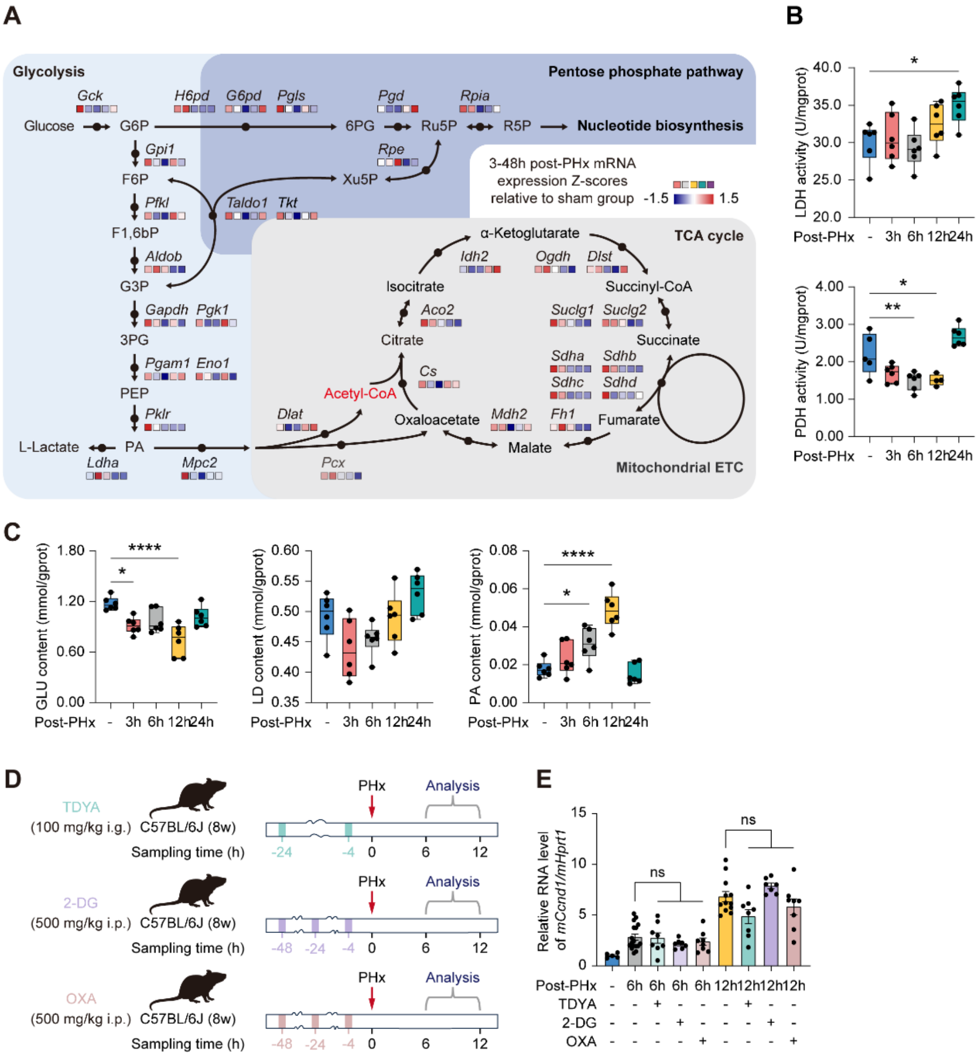
A
Glycolysis
Pentose phosphate pathway
Gck
H6pd
G6pd
Pgls
Pgd
Rpia
Glucose
G6P
6PG
Ru5P
R5P
Nucleotide biosynthesis
Gpi1
Rpe
F6P
Xu5P
3-48h post-PHx mRNA expression Z-scores relative to sham group
-1.5
1.5
Pfkl
Taldo1
Tkt
F1,6bP
α-Ketoglutarate
TCA cycle
Aldob
Idh2
Ogdh
Dlst
G3P
Isocitrate
Succinyl-CoA
Gapdh
Pgk1
Aco2
Suclg1
Suclg2
3PG
Citrate
Succinate
Pgam1
Eno1
Cs
Sdha
Sdhb
Sdhc
Sdhd
PEP
Acetyl-CoA
Pklr
Dlat
Oxaloacetate
Mdh2
Fh1
Fumarate
L-Lactate
PA
Malate
Ldha
Mpc2
Pcx
Mitochondrial ETC
B
LDH activity (U/mgprot)
PDH activity (U/mgprot)
Post-PHx
-
3h
6h
12h
24h
C
GLU content (mmol/gprot)
LD content (mmol/gprot)
PA content (mmol/gprot)
D
TDYA
(100 mg/kg i.g.)
C57BL/6J (8w)
2-DG
(500 mg/kg i.p.)
OXA
(500 mg/kg i.p.)
Sampling time (h)
PHx
Analysis
E
Relative RNA level of mCcnd1/mHprt1
ns
Post-PHx
TDYA
2-DG
OXA

**Fig. S7. Changes in the central carbon metabolic network in the initiation phase of liver regeneration.**

(A) Glycolysis, tricarboxylic acid cycle (TCA), and oxidative phosphorylation pathway. These pathways are interconnected and constantly in flux. FPKM at different time points from 3 to 48 hours were normalized relative to FPKM of the sham group. Z-scores were visualized with color ranging from red (high expression relative to sham) over white (intermediate expression relative to sham) to blue (low expression relative to sham).

(B) Comparison of hepatic lactate dehydrogenase (LDH) activity and pyruvate dehydrogenase (PDH) activity in regenerating livers at different time. Data are represented as mean ± SEM of 6 replicates.

(C) Comparison of hepatic glucose (GLU) content, lactic acid (LD) content, and pyruvic acid (PA) content in regenerating livers at different time. Data are represented as mean ± SEM of 6 replicates.

(D) Animal experimental strategy of inhibitor treatment.

(E) qRT-PCR analysis of the expression of *Ccnd1* mRNA. Data are represented as mean ± SEM of 7 to 12 replicates.

ns, not significant, $^{*}P < 0.05$; $^{**}P < 0.01$; $^{***}P < 0.001$; $^{****}P < 0.0001$, all compared with control. Error bars represent mean ± SEM.

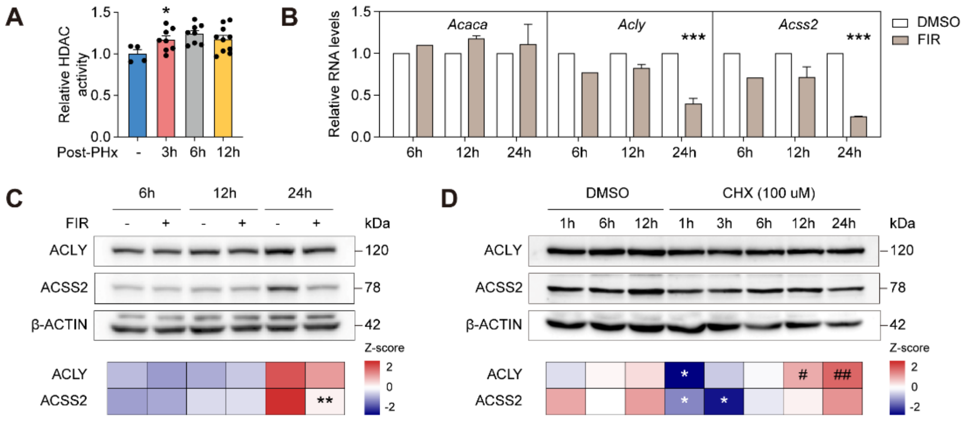
A
Relative HDAC activity
Post-PHx
3h
6h
12h
B
Relative RNA levels
Acaca
Acly
Acss2
DMSO
FIR
C
ACLY
ACSS2
β-ACTIN
kDa
Z-score
D
CHX (100 uM)

371

372

**Fig. S8. The expression of enzymes related to Acetyl-CoA production.**

(A) Hepatic HDAC activity.

(B) mRNA levels of *Acaca*, *Acly*, and *Acss2* in HepG2 cells treated with or without ACC inhibitor, FIR (300 nM). $^{***}P < 0.001$; $^{****}P < 0.0001$, all compared with DMSO control at the same time point.

(C) Western blot analysis and quantification of enzymes involved in acetyl-CoA production in HepG2 cells with or without FIR treatment (300 nM). $^{**}P < 0.01$, compared with DMSO control at the same time point.

(D) Western Blot analysis and quantification of enzymes involved in acetyl-CoA production in HepG2 cells treated with CHX (100 mM) at indicated time points. $^{*}P < 0.05$, compared with DMSO control at 1 hour. $^{\#}P < 0.05$; $^{\#\#}P < 0.01$, compared with CHX treatment at 1 hour.

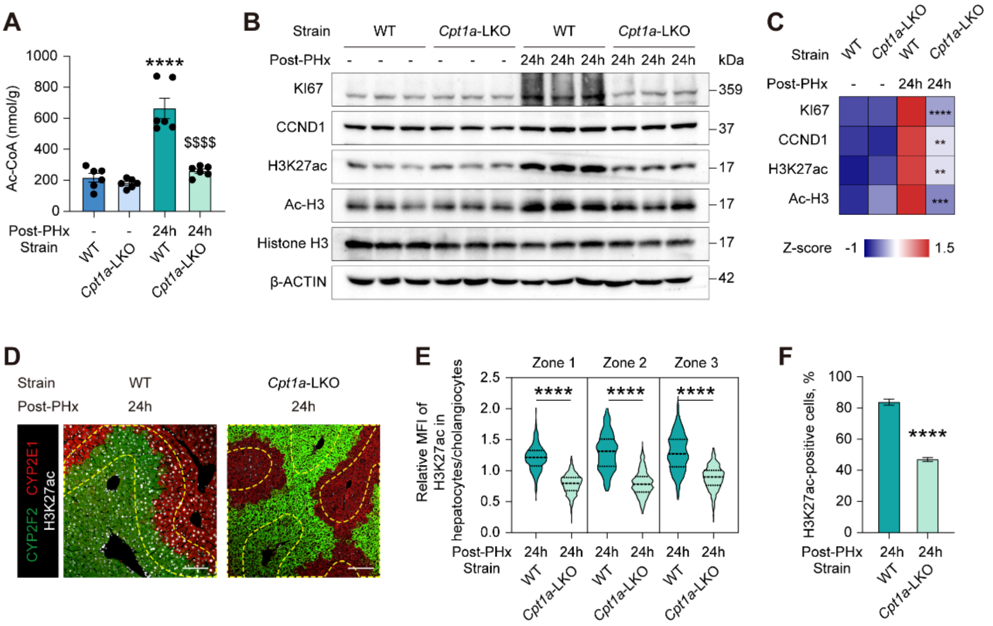
A
Ac-CoA (nmol/g)
1000
800
600
400
200
0
****
$$$$
Post-PHx - - 24h 24h
Strain WT Cpt1a-LKO WT Cpt1a-LKO
B
Strain WT Cpt1a-LKO WT Cpt1a-LKO
Post-PHx - - - - - - 24h 24h 24h 24h 24h 24h
kDa
KI67 359
CCND1 37
H3K27ac 17
Ac-H3 17
Histone H3 17
β-ACTIN 42
C
Strain WT Cpt1a-LKO WT Cpt1a-LKO
Post-PHx - - 24h 24h
KI67 ****
CCND1 **
H3K27ac **
Ac-H3 ***
Z-score -1 1.5
D
Strain WT Cpt1a-LKO
Post-PHx 24h 24h
CYP2F2 CYP2E1 H3K27ac
E
Relative MFI of H3K27ac in hepatocytes/cholangiocytes
Zone 1 Zone 2 Zone 3
2.5
2.0
1.5
1.0
0.5
0.0
**** **** ****
Post-PHx 24h 24h 24h 24h 24h 24h
Strain WT Cpt1a-LKO WT Cpt1a-LKO WT Cpt1a-LKO
F
H3K27ac-positive cells, %
100
80
60
40
20
0
****
Post-PHx 24h 24h
Strain WT Cpt1a-LKO

**Fig. S9. The hepatic CPT1A deletion inhibits histone acetylation and cell cycle progression after PHx.**

(A) Comparison of hepatic Acetyl-CoA levels in WT and *Cpt1a*-LKO mice before and 24 hours after PHx.

(B and C) Western blotting analysis and quantitation of proliferative hallmarks and histone acetylation modifications in livers of WT and *Cpt1a*-LKO mice before and 24 hours after PHx. H3 was used as a loading control. Histone H3 is used as loading control for acetylation modifications. β-ACTIN serves as loading control for other proteins.

(D-F) Immunofluorescence staining for CYP2E1, CYP2F2, and H3K27ac on sections from regenerating livers of WT and *Cpt1a*-LKO mice at 24 hours after PHx (G). The relative mean fluorescence intensity (MFI) of H3K27ac in hepatocytes compared to cholangiocytes across different liver zonation was represented in H. Proportion of H3K27ac$^{+}$ hepatocytes across sections is in I. Scale bar, 100 μm. CYP2E1$^{+}$ cells in red represent the central vein area, while CYP2F2$^{+}$ cells in green represent the portal vein area. The yellow dashed lines indicate Zone 2 lobule. Quantifications were performed on at least 5 different areas of the sections in a random way.

$^{*}P<0.05$; $^{**}P<0.01$; $^{***}P<0.001$; $^{****}P<0.0001$, unpaired Student's t test. Error bars represent mean ± SEM.

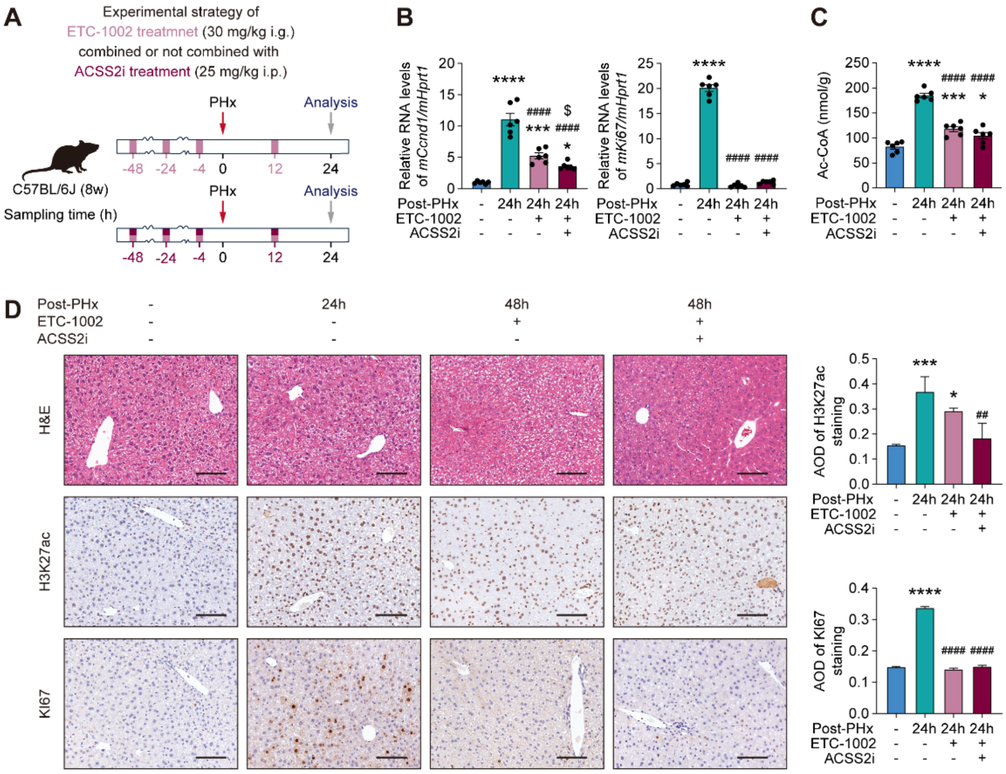
A
Experimental strategy of
ETC-1002 treatmnet (30 mg/kg i.g.)
combined or not combined with
ACSS2i treatment (25 mg/kg i.p.)
C57BL/6J (8w)
Sampling time (h)
PHx
Analysis
-48 -24 -4 0 12 24
B
Relative RNA levels of mCcnd1/mHprt1
Relative RNA levels of mKi67/mHprt1
Post-PHx
ETC-1002
ACSS2i
C
Ac-CoA (nmol/g)
D
Post-PHx - 24h 48h 48h
ETC-1002 - - + +
ACSS2i - - - +
H&E
H3K27ac
KI67
AOD of H3K27ac staining
AOD of KI67 staining

**Fig. S10. Histone acetylation and the initiation of liver regeneration is dependent on the production of Acetyl-CoA.**

(A) Animal experimental design for the treatment of ACLY and ACSS2 inhibitors.

(B) qRT-PCR analysis of the mRNA levels of *Ccnd1* and *Ki67*. Data are represented as mean ± SEM of 6 replicates.

(C) Hepatic Acetyl-CoA contents.

(D) Representative images of H&E and immunohistochemical staining for regenerating liver with or without ACLY and ACSS2 inhibitors, with quantitative analysis of the AOD value. Scale bars, 100 μm. Quantifications were performed on at least 5 different areas of the sections in a random way.

ns, not significant, $^{*}P < 0.05$; $^{**}P < 0.01$; $^{***}P < 0.001$; $^{****}P < 0.0001$, Error bars represent mean ± SEM.

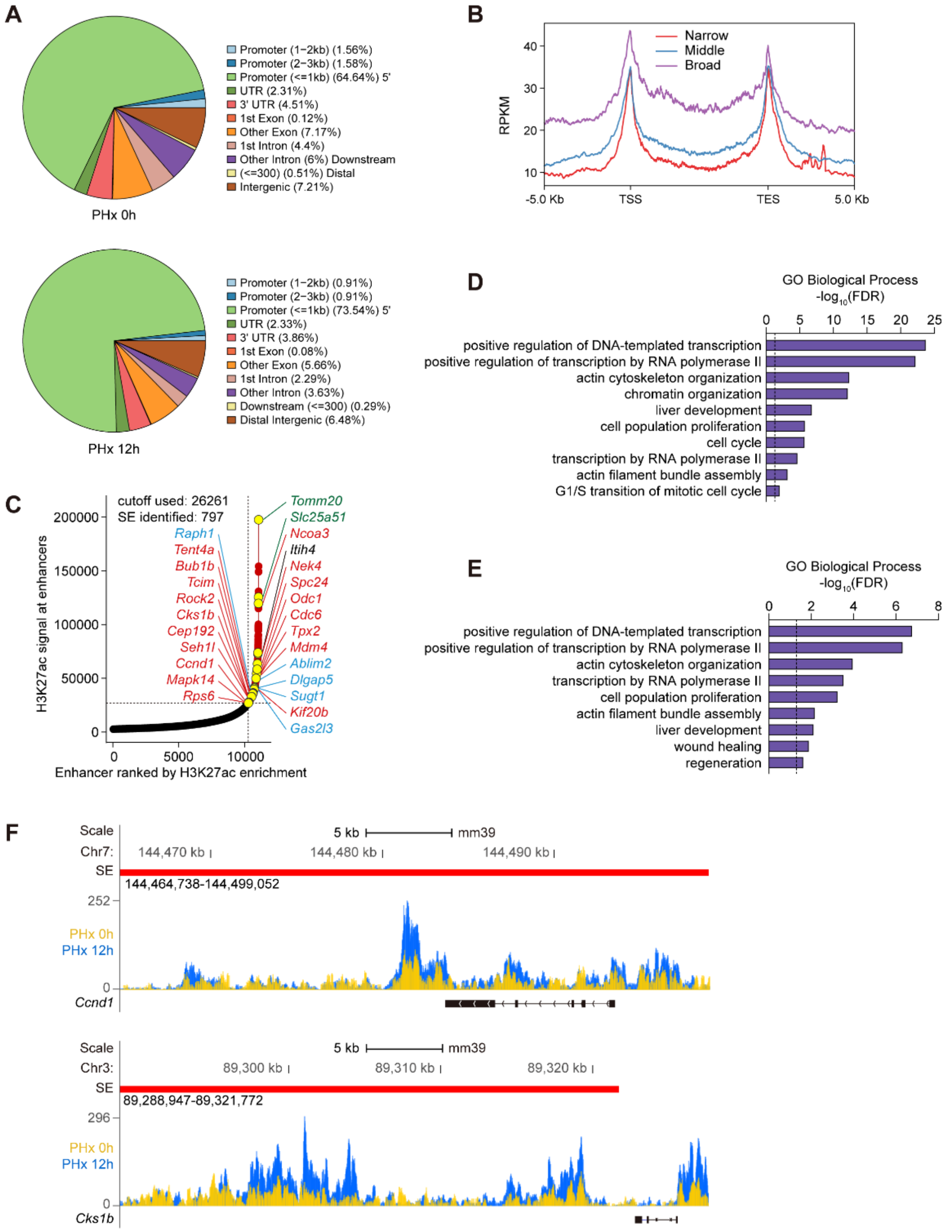
A
Promoter (1−2kb) (1.56%)
Promoter (2−3kb) (1.58%)
Promoter (<=1kb) (64.64%) 5'
UTR (2.31%)
3' UTR (4.51%)
1st Exon (0.12%)
Other Exon (7.17%)
1st Intron (4.4%)
Other Intron (6%) Downstream
(<=300) (0.51%) Distal
Intergenic (7.21%)
PHx 0h
Promoter (1−2kb) (0.91%)
Promoter (2−3kb) (0.91%)
Promoter (<=1kb) (73.54%) 5'
UTR (2.33%)
3' UTR (3.86%)
1st Exon (0.08%)
Other Exon (5.66%)
1st Intron (2.29%)
Other Intron (3.63%)
Downstream (<=300) (0.29%)
Distal Intergenic (6.48%)
PHx 12h
B
Narrow
Middle
Broad
RPKM
-5.0 Kb
TSS
TES
5.0 Kb
C
cutoff used: 26261
SE identified: 797
H3K27ac signal at enhancers
Enhancer ranked by H3K27ac enrichment
Tomm20
Slc25a51
Ncoa3
Itih4
Nek4
Spc24
Odc1
Cdc6
Tpx2
Mdm4
Ablim2
Dlgap5
Sugt1
Kif20b
Gas2l3
Raph1
Tent4a
Bub1b
Tcim
Rock2
Cks1b
Cep192
Seh1l
Ccnd1
Mapk14
Rps6
D
GO Biological Process
-log10(FDR)
positive regulation of DNA-templated transcription
positive regulation of transcription by RNA polymerase II
actin cytoskeleton organization
chromatin organization
liver development
cell population proliferation
cell cycle
transcription by RNA polymerase II
actin filament bundle assembly
G1/S transition of mitotic cell cycle
E
GO Biological Process
-log10(FDR)
positive regulation of DNA-templated transcription
positive regulation of transcription by RNA polymerase II
actin cytoskeleton organization
transcription by RNA polymerase II
cell population proliferation
actin filament bundle assembly
liver development
wound healing
regeneration
F
Scale
Chr7:
SE
5 kb
mm39
144,470 kb
144,480 kb
144,490 kb
144,464,738-144,499,052
252
PHx 0h
PHx 12h
Ccnd1
Scale
Chr3:
SE
5 kb
mm39
89,300 kb
89,310 kb
89,320 kb
89,288,947-89,321,772
296
PHx 0h
PHx 12h
Cks1b

**Fig. S11. CUT&Tag sequencing analysis for genes directly regulated by H3K27ac modification after PHx.**

(A) H3K27ac CUT&Tag peaks classified by mouse genomic annotations (mm9) in PHx 0h and PHx 12h.

(B) Coverage plots of H3K27ac CUT&Tag signals within genes in three different groups categorized according to the breadth of H3K27ac peaks in PHx 12h group. The 25% of genes with the broadest peaks were placed in the 'broad' group, the 25% of genes with the narrowest peaks were placed in the 'narrow' group, and the 'medium' group contains all other genes (25%–75%). RPKM, reads per million mapped reads; TSS, transcription start site; TES, transcription end site.

(C) Call of super-enhancers (SEs) with H3K27ac CUT&Tag data using ROSE algorithm analysis. The distribution of H3K27ac tag intensities revealed the locations of predicted 797 SEs showing unevenly high H3K27ac signals in red dots, while the rest black dots represent typical enhancers. Known liver regeneration-associated genes proximal to predicted SEs are indicated in yellow dots, including mitochondrial respiration-related (green annotation), cytoskeleton-related (blue annotation), and cell cycle-related (red annotation) genes.

(D and E) Top significantly associated biological processes associated with distal regulatory elements (enhancers in upper panel and predicted SEs in lower panel).

(E) Genome browser tracks of H3K27ac CUT&Tag signals at the *Ccnd1* (upper panel) and *Cks1b* (lower panel) locus showed stretches of predicted enhancers, corresponding to predicted SEs with high H3K27ac signals.

### Supplementary tables

**Table S1. Identification of differential H3K27ac peaks between PHx 12h and PHx 0h.**

**Table S2. Up-target and down-target genes in PHx 12h group predicted by BETA algorithms.**

**Table S3. The transcriptional regulatory networks of H3K27ac-related genes predicted from the ChEA3 database.**

**Table S4. Key Resource Table.**

| REAGENT or RESOURCE | SOURCE | IDENTIFIER |
|---|---|---|
| **Antibodies** | | |
| PCNA (PC10) Mouse mAb | Cell Signaling Technology | Cat #: 2586s; RRID: AB_2160343 |
| KI67 Polyclonal antibody | proteintech | Cat #: 27309-1-AP; RRID: AB_2756525 |
| Beta Actin Monoclonal antibody | proteintech | Cat #: 66009-1-lg; RRID: AB_2687938 |
| Recombinant Anti-Cyclin D1 antibody (EPR2241) - C-terminal | Abcam | Cat #: ab134175; RRID: AB_2750906 |
| Cyclin D1 Monoclonal antibody | proteintech | Cat #: 60186-1-lg; RRID: AB_10793718 |
| Acetyl-Histone H3 (Lys27) Antibody | Cell Signaling Technology | Cat #: 4353S; RRID: AB_10545273 |
| Recombinant Anti-Histone H3 (acetyl K27) antibody (EP16602) - ChIP Grade | Abcam | Cat #: ab177178; RRID: AB_2828007 |
| Histone-H3 Polyclonal antibody | proteintech | Cat #: 17168-1-AP; RRID: AB_2716755 |
| Ac-Histone H3 Antibody (AH3-120) | Santa Cruz | Cat #: sc-56616; RRID: AB_2263811 |
| Ac-lysine Antibody (AKL5C1) | SantaCruz | Cat #: sc-32268; RRID: AB_627898 |
| ACSS2 Antibody (A-9) | SantaCruz | Cat #: sc-398559; RRID: AB_3086604 |
| ATP-citrate synthase Antibody (5F8D11) | SantaCruz | Cat #: sc-517267; RRID: AB_3086605 |
| CYP2E1-Specific Polyclonal antibody | proteintech | Cat #: 19937-1-AP; RRID: AB_10646444 |
| CYP2F2 Antibody (F-9) | SantaCruz | Cat #: sc-374540; RRID: AB_10987684 |

| | | |
|---|---|---|
| HRP-conjugated Affinipure Goat Anti-Mouse IgG(H+L) | proteintech | Cat #: SA00001-1; RRID: AB_2722565 |
| Goat anti-Rabbit IgG-HRP Antibody | absin | Cat #: abs20040; RRID: N/A |
| **Chemicals, peptides, and recombinant proteins** | | |
| (R)-Etomoxir sodium salt (ANTI-EAPII (TTRAP)) | Shanghai yuanye | Cat #: S80976; CAS: 828934-41-4 |
| 10,12-Tricosadiynoic Acid | Aladdin | Cat #: T162085; CAS: 66990-30-5 |
| 2-Deoxy-D-glucose | Shanghai yuanye | Cat #: S11070; CAS: 154-17-6 |
| sodium oxamate | Shanghai yuanye | Cat #: S30701; CAS: 565-73-1 |
| C646 | Topscience | Cat #: T2452; CAS: 328968-36-1 |
| Firsocostat | Topscience | Cat #: T7184; CAS: 1434635-54-7 |
| Cycloheximide | Aladdin | Cat #: C112766; CAS: 66-81-9 |
| DMEM medium | VivaCell | Cat #: C3113-0500; Lot No.: 2315078 |
| Penicillin-Streptomycin Solution | KeyGEN | Cat #: KGL2303-100; Lot No.: 20220519 |
| Fetal Bovine Serum | VivaCell | Cat #: C2910-0500; Lot No.: 2038153 |
| RNA isolater Total RNA Extraction Reagent | Vazyme | Cat #: R401-01 |
| HiScript III RT SuperMix for qPCR (+gDNA wiper) | Vazyme | Cat #: R323-01 |
| Taq Pro Universal SYBR qPCR Master Mix | Vazyme | Cat #: Q712-02 |
| RIPA Lysis Buffer | Beijing Aoqing | Cat #: AQ522 |
| Protease Inhibitor Cocktail | Beijing Aoqing | Cat #: AQ551 |
| SDS-PAGE loading buffer,4X (with DTT) | Biorigin | Cat #: BN20104 |
| Color Prestained Protein Marker | LABLEAD | Cat #: P1018 |
| Mounting Medium With DAPI - Aqueous, Fluoroshield | Abcam | Cat #: ab104139 |
| **Critical commercial assays** | | |
| Acetyl CoA Content Assay Kit | Solarbio | Cat #: BC0980 |
| Histone Acetyltransferase Activity Assay Kit (Colorimetric) | Abcam | Cat #: ab65352 |
| HDAC Assay Kit (Colorimetric) | ACTIVE MOTIF | Cat #: 56210 |
| IHC Detect Kit for Rabbit/Mouse Primary Antibody | proteintech | Cat #: PK10006 |
| Five-color Fluorescence kit | Recordbio | Cat #: RC 0086-45 |
| BCA Protein Assay Kit | Biorigin | Cat #: BN27109 |
| Super ECL Plus | Biorigin | Cat #: BN16009 |
| Total cholesterol assay kit | Nanjingjiancheng | Cat #: A111-1-1 |
| Lactate dehydrogenase assay kit | Nanjingjiancheng | Cat #: A020-2-2 |
| Pyruvate assay kit | Nanjingjiancheng | Cat #: A081-1-1 |

| | | |
|---|---|---|
| Pyruvate Dehydrogenase (PDH) Activity Assay Kit | Solarbio | Cat #: BC0380 |
| Lactic Acid assay kit | Nanjingjiancheng | Cat #: A019-2-1 |
| Glucose kit (glucose oxidase method) | Nanjingjiancheng | Cat #: A154-1-1 |
| **Experimental models: Cell lines** | | |
| Mouse: HepG2 cells | This paper | N/A |
| **Experimental models: Organisms/strains** | | |
| Mouse: C57/BL6J | SPF Biotechnology Co., Ltd. (Beijing, China) | N/A |
| Mouse: *Cpt1a*$^{fl/fl}$ Alb-cre mice | GemPharmatech Co., Ltd (Nanjing, China) | N/A |
| **Primers** | | |
| *mCcnd1* Forward 5'-CGTATCTTACTTCAAGTGCGTG-3' | Sangon Biotech (Shanghai) Co., Ltd. | N/A |
| *mCcnd1* Reverse 5'-ATGGTCTCCTTCATCTTAGAGG-3' | Sangon Biotech (Shanghai) Co., Ltd. | N/A |
| *mKi67* Forward 5'-CCTGTCACTCCAGATCAGAACT-3' | Sangon Biotech (Shanghai) Co., Ltd. | N/A |
| *mKi67* Reverse 5'-GGAGGCAGTCTTCATAGTCTTCT-3' | Sangon Biotech (Shanghai) Co., Ltd. | N/A |
| *mPcna* Forward 5'-CCGAGACCTTAGCCACATTG-3' | Sangon Biotech (Shanghai) Co., Ltd. | N/A |
| *mPcna* Reverse 5'-CCTCCTCTTCTTTATCCACATTAC-3' | Sangon Biotech (Shanghai) Co., Ltd. | N/A |
| *mHprt1* Forward 5'-CCGAGGATTTGGAAAAAGTGTT-3' | Sangon Biotech (Shanghai) Co., Ltd. | N/A |
| *mHprt1* Reverse 5'-CATCTCCTTCATGACATCTCGA-3' | Sangon Biotech (Shanghai) Co., Ltd. | N/A |
| *mCpt1a* Forward 5'-CTACATCACCCCAACCCATATT-3' | Sangon Biotech (Shanghai) Co., Ltd. | N/A |

| | | |
|---|---|---|
| *mCpt1a* Reverse 5’-GATCCCAGAAGACGAATAGGTT-3’ | Sangon Biotech (Shanghai) Co., Ltd. | N/A |
| *hAcaca* Forward 5’-TCTCCTCCAACCTCAACCACTATG-3’ | Sangon Biotech (Shanghai) Co., Ltd. | N/A |
| *hAcaca* Reverse 5’-ATTCCGCCCATCCGCTGAC-3’ | Sangon Biotech (Shanghai) Co., Ltd. | N/A |
| *hAcly* Forward 5’-CAACTTTGCCTCTCTCCGCTCTG-3’ | Sangon Biotech (Shanghai) Co., Ltd. | N/A |
| *hAcly* Reverse 5’-TCCCTTCTGGTCCGCCTTCTTG-3’ | Sangon Biotech (Shanghai) Co., Ltd. | N/A |
| *hAcss2* Forward 5’-CCATTGCCACACCAGACTACATCC-3’ | Sangon Biotech (Shanghai) Co., Ltd. | N/A |
| *hAcss2* Reverse 5’-CTTCCGAAGCACTCGCCTCATG-3’ | Sangon Biotech (Shanghai) Co., Ltd. | N/A |
| *hHprt1* Forward 5’-TATGGCGACCCGCAGCCCT-3’ | Sangon Biotech (Shanghai) Co., Ltd. | N/A |
| *hHprt1* Reverse 5’-CATCTCGAGCAAGACGTTCAG-3’ | Sangon Biotech (Shanghai) Co., Ltd. | N/A |
